# Distinct Orbitofrontal Feedback Signals Shape Sensory Behavioral Strategies during Flexible Learning

**DOI:** 10.64898/2026.07.15.737076

**Authors:** Jasper Teutsch, Rohan Rao, Mark D. Humphries, Silvia Maggi, Abhishek Banerjee

## Abstract

Animals adapt their behavior by integrating sensory evidence with prior experience and contextual information. Understanding how animals employ specific behavioral strategies to integrate these variables and how neural mechanisms support such strategies remains unclear. We trained mice on a tactile reversal learning task and used a trial-by-trial computational model to infer latent decision strategies. Early in learning, mice relied on action-based (“choice-driven”) policies but progressively transitioned to stimulus-guided choices (“cue-driven”) as they learned the task. Following a rule reversal, mice flexibly reinstated this policy to adapt their behavior. Chemogenetic silencing of the lateral orbitofrontal cortex (lOFC) delayed this transition and impaired reversal learning. To identify the neural basis of strategy learning, we developed a novel approach combining low-dimensional analysis of neural activity with decoding of behavioral strategies from longitudinal two-photon imaging. We revealed reward- and error-strategy representations in excitatory layer 2/3 neurons in the primary somatosensory cortex (S1) that were differentially modulated by OFC instructive signals. Within S1, distinct subpopulations of positive and negative-valence-coding neurons tracked evolving decision strategies through trial-history integration. Together, these findings revealed how mice flexibly deploy distinct exploratory strategies during adaptive behavior, highlighting OFC’s role in supporting reward and error-guided learning through distinct corticocortical interactions.

## Introduction

Animals adapt their behavior by integrating sensory evidence with prior experience and contextual information to optimize their decisions in novel environments. At a mechanistic level, this adaptability depends on causally linking sensory inputs and actions to rewarding or aversive outcomes, enabling efficient exploration and rapid updating of behavior^1^. Such reward- and error-driven learning underlies behavioral flexibility and is disrupted across multiple neurological disorders^2^. However, how distinct behavioral strategies emerge during learning and are implemented across distributed neural circuits remains poorly understood.

Frontal cortical areas are central to executive function, supporting rule-based strategies that guide task performance across species. Positioned at the apex of a cortical processing hierarchy, these regions encode task-relevant variables that shape behavior across contexts^3–4^. Within this framework, the orbitofrontal cortex (OFC) has been shown to play a critical role in constructing and updating stimulus–outcome and action–outcome associations, and in tracking task structure across trials^5–7^. A prominent view proposes that the OFC functions as a cognitive map of task space^8^, representing predictions about potential outcomes and their probabilities based on past and current choices. These representations are thought to provide a substrate for flexible readout by downstream circuits to support learning and behavioral adaptation^9–10^. Consistent with this view, perturbing OFC function induces behavioral perseveration and impaired contingency updating^11^, while simultaneously weakening task-variable representations across distributed cortical and subcortical regions. Recent work further demonstrates that feedback from lateral OFC (lOFC) reshapes non-sensory representations in the primary somatosensory cortex (S1) during contingency switches^9,12^, raising the question of whether sensory cortices actively participate in adaptive computations rather than passively reflecting top-down signals.

Accumulating evidence supports this view, revealing that neurons in S1 actively encode not only sensory features but also abstract, task-related variables, including reward reinforcement^13^, decision variables^14^, the integration of outcome-history^9^, and surprise or prediction errors^15^. These key variables are central to adopting and updating specific behavioral strategies, yet how they are integrated to guide trial-by-trial behavioral adaptation is unclear. Importantly, such strategy-related representations may emerge at the level of population dynamics whilst also being reflected in the heterogeneous activity of individual neurons. Consistent with this idea, population-level neural responses in S1 reorganize over learning^14,16^, suggesting that neuronal ensembles encode decision-related task variables that evolve over learning. These observations suggest that S1 could function as a dynamic computational hub integrating sensory, contextual, and outcome-related information. Yet, precisely how contextual learning recruits diverse S1 subpopulations to support behavioral strategies, and whether such representations depend on frontal feedback, remains largely unanswered.

Here, we address these questions by combining a tactile reversal learning paradigm with longitudinal two-photon calcium imaging and trial-by-trial computational inference of behavior. Using a Bayesian analysis framework, we inferred latent reward- and error-based strategies guiding decision-making. To link behavior to neural dynamics, we developed a novel approach that aligns low-dimensional population activity with strategy decoding at single-trial resolution. We show that mice transition from action-based to stimulus-guided strategies during learning and reinstated this policy following rule reversal. Chemogenetic silencing of lOFC disrupts these transitions, prolongs reliance on recent trial history, and impairs adaptive updating during reversal. At the neural level, we identify distinct excitatory subpopulations in S1 that selectively encoded reward- and error-based behavioral strategies. Population-level analyses reveal that strategy-related signals are contained in a latent subspace that dynamically track behavioral strategy use across learning. Additionally, we identified a single-neuron mechanism that integrated past and current trial information within a specialized outcome-selective S1 subpopulation. Critically, both population-level strategy representation and single-neuron integration of strategy-specific outcome were dependent on lOFC input, demonstrating a causal role for distinct frontal feedback signals in shaping sensory representations during adaptive behavior. Taken together, these findings reveal S1 as a key cortical locus where somatosensory information, recent experience-history, and top-down frontal inputs converge to support the flexible, adaptive use of behavioral strategies.

## Results

### Choice-to-cue strategy switch drives flexible learning

To investigate how animals use distinct behavioral strategies during learning, we trained mice in a tactile reversal learning task based on a ‘Go/No-Go’ paradigm^9^. Mice initially learned to discriminate between two distinct sandpaper textures (coarse P100 and fine P1200) by either licking for the ‘Go’ (P100) stimulus to win a water reward or withholding licking for the ‘No-Go’ (P1200) stimulus to avoid a mild white noise (**Fig. 1a**; see **Methods**). All mice successfully learnt the task (from ‘learning naïve’, LN, through ‘learning expert’, LE). Upon expert performance (*d*’ > 1.5), the stimulus-outcome contingencies were reversed (‘reversal learning’). Following rule reversal, the performance of all mice initially decreased in the ‘reversal naïve’ (RN) phase but subsequently reached expert levels (‘reversal expert’, RE) (**Fig. 1b**). To study how animals dynamically applied explorative strategies during flexible learning, we implemented a Bayesian evidence accumulation model^17^. Bayesian analysis enabled the tracking, on a trial-by-trial basis, of the probability that animals used a specific behavioral strategy (P(strategy)) (**Fig. 1c**; see **Methods**). Strategy probabilities were calculated by comparing three task variables (stimulus (S), action (A), and outcome (O)) across consecutive trial pairs (e.g., *t-1* and *t*) during learning. We defined ‘cue-based’ strategies as those contingent on stimulus–outcome (S–O) associations, and ‘choice-based’ strategies as those contingent on action–outcome (A–O) associations (**Fig. 1d**).

**Figure 1.**
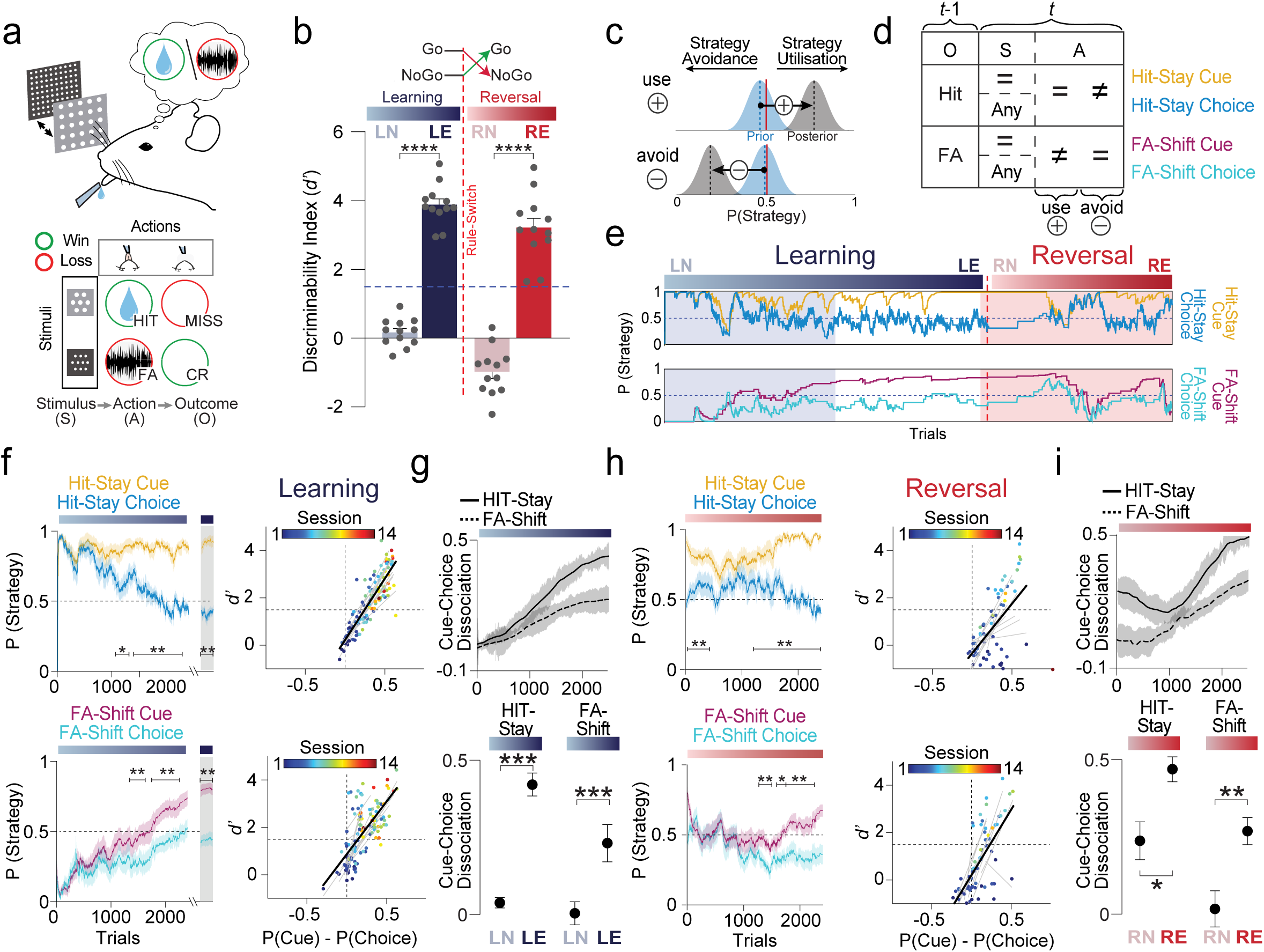
Behavioral adaptation upon reversal learning follows a choice-to-cue strategy transition. **a**, Top, schematic of the experimental setup for the head-fixed Go/No-Go texture discrimination task. Upon presentation of one of two stimuli (S), mice performed an action (A) leading to outcomes (O) such as a reward (hit), a punishment (false alarm, FA) or an omission of both (miss or correct rejection (CR)). Bottom, schematic of trial-outcome types (before rule-switch) with two different ‘stimulus-action’ (S-A) combinations resulting in ‘win’ (green circles, hit or CR) or ‘loss’ (red circles, FA or miss), respectively. **b**, Once mice trained on the tactile discrimination task displayed expert performance during learning, stimulus-outcome contingencies were reversed. Task performance was measured as the discriminability index (*d*’) of WT mice (*n* = 12) at four salient task phases (LN, learning naïve; LE, learning expert; RN, reversal naïve; RE, reversal expert). **c**, Bayesian analysis of exploratory strategies based on trial-history-dependent task parameters. Based on the combination of a strategy-specific set of parameters, the model updates the prior in trial *t-1* to the posterior distribution either positively (strategy implementation) or negatively (strategy avoidance). The posterior is then used as the new prior in the subsequent trial comparison. If trial *t-1* was a ‘hit’ and the stimulus presented in both consecutive trials was the same, a repetition of the same action (‘stay’) would result in a positive update of the ‘hit-stay cue’ strategy prior, while a change of action (‘shift’) would cause a negative update. **d**, An overview of parameters defining each strategy. For cue-based strategies, stimulus (*t-1*, *t*), action (*t-1*, *t*), and outcome (*t-1*) parameters were considered, while choice-based strategies were independent of stimulus presentation. **e**, Dynamics of strategy implementation (P(strategy)) for one example mouse during learning and reversal learning. **f**, Left, comparison of cue and choice-based strategy dynamics during initial learning and expert performance (LE) for ‘hit-stay’ and ‘FA-shift’ strategies. The curves show the mean ± S.E.M. over trials. Right, scatterplots displaying the relationship of d’ and cue-choice dissociation (P(cue)-P(choice)) for the respective strategies (‘hit-stay’, ‘FA-shift’) in each session. Each data point is one individual session and the color indicates the session number. Grey lines indicate linear fits for individual animals; the average of all individual fits is shown as the black line. **g**, Dynamics and quantification of the cue-choice dissociation during the initial learning phase. **h**, same as **f**, but for reversal learning. **i**, same as **g**, but for reversal learning. n = 12 WT mice for all plots. Data presented as mean ± S.E.M.; *p<0.05, **p<0.01, ***p<0.001, paired two tailed t-test. Horizontal bars indicate significant differences between curves at the corresponding trials, as determined by a cluster-based permutation test.

To specifically examine strategies guiding reward- and error-based learning, the process by which animals potentially link stimuli, actions and contexts to rewards and losses, we analysed variants of commonly described ‘win-stay’ (repeat choice after a win) and ‘lose-shift’ (choose alternatives after a loss) strategies^18–20^. By focusing on unambiguously rewarded and punished outcomes (hit = reward, FA = punishment), we quantified four strategies: ‘hit-stay cue’, ‘hit-stay choice’, ‘FA-shift cue’, and ‘FA-shift choice’, to dissociate S-O and A-O associations that are crucial for task learning^21^ (**Fig. 1d,e**). During initial learning (LN→LE), we observed a gradual dissociation of ‘cue’ and ‘choice’-based strategies (‘cue-choice dissociation’) in both ‘hit-stay’ and ‘FA-shift’ that strongly correlated with task performance (**Fig. 1f,g**). The relationship of cue-choice dissociation and stimulus-driven behavior can be further illustrated using simulated data (**Supplementary Fig. 1a,b**) in which transitions from action-based (A–O) to stimulus-based (S–O) choice policies resulted in increasing dissociation values. Incorporating mice’s lick bias into the simulations reproduced the differential trajectories observed for ‘hit-stay’ and ‘FA-shift’ strategies during initial learning (LN) (**Supplementary Fig. 1c,d**). Notably, the speed and asymptotic magnitude of dissociation differed between strategies, with ‘FA-shift’ exhibiting a slower and smaller increase than ‘hit-stay’ (**Fig. 1f,g**).

Following a rule-switch, ‘cue’ and ‘choice’-based strategies rapidly reconverged before gradually dissociating as performance improved (RN→RE) (**Fig. 1h,i**). This reconvergence reflected a transient shift in favor of action-driven choices immediately after reversal. The partial cue-choice reconvergence observed for ‘hit-stay’ following the rule-switch was primarily attributable to transient disengagement in a subset of animals (**Supplementary Fig. 1d**), resulting in fewer trials meeting strategy criteria. As during initial learning, mice subsequently showed a slower and smaller cue-choice dissociation in ‘FA-shift’ compared to ‘hit-stay’ in the expert reversal phase (RE).

Analysis of individual differences revealed that naïve animals used similar strategies (LN) (**Supplementary Fig. 1e,f**), driven in part by a global bias towards licking across animals (**Supplementary Fig. 1d**). As animals transitioned to expert performance (LN→LE), inter-individual variability increased, and animals showed a relative bias toward reward-driven over error-driven strategies (positive ΔHit-Stay/FA-Shift) (**Supplementary Fig. 1g**). Interestingly, individual variability peaked immediately after a rule-switch (RN), with some animals exhibiting a transient shift towards error-driven strategies (negative ΔHit-Stay/FA-Shift). When transitioning to the expert phase (RE), animals’ behavior reconverged with variability returning to LE levels, and all animals relied more on reward-driven learning (**Supplementary Fig. 1e-g**). Collectively, these results reveal that mice exhibited consistent population-level behavioral dynamics during learning and reversal, while individual animals differ in the timing and extent of strategy transitions as choices transitioned from action- to stimulus-driven for both reward- and error-based learning. Notably, the onset of animals’ cue-choice dissociation occurred closely around the ‘learning point’, further highlighting cue-choice dissociation’s close relationship to the improvement of general task learning performance (**Supplementary Fig. 1h,i**).

### Lateral OFC silencing differentially impairs reward- and error-driven strategies during reversal learning

Previous work has highlighted OFC’s key role in credit assignment and outcome-based reward learning, especially when previously learnt S-O associations are altered and require updating^18,22–24^. To examine the contribution of the lOFC to strategy dynamics during reversal learning, we chemogenetically silenced excitatory layer (L)2/3 neurons in the lOFC by locally expressing inhibitory DREADD receptors (hM4Di) (**Fig. 2a**). Systemic injection of CNO, activating hM4Di and thereby silencing neurons following rule-switch, significantly impaired behavioral adaptation during reversal learning (**Fig. 2b**). Only a subset (n = 4 out of 6) of lOFC-silenced mice reached expert performance, and those animals required significantly more trials to do so. The reversal rate of this subset was significantly delayed, as evidenced by increased trial-to-criterion (**Fig. 2b**) and delayed learning points (**Supplementary Fig. 2a**). In contrast to WT controls, lOFC-silenced mice exhibited reduced cue-choice dissociation for both reward- and error-driven strategies during reversal learning (**Fig. 2c,d**). While the emergence of cue-choice dissociation was delayed for both ‘hit-stay’ and ‘FA-shift’ strategies, a pronounced reduction in the asymptotic magnitude of dissociation was observed specifically for reward-driven (‘hit-stay’) strategies (**Fig. 2c,d**). When cue–choice dissociation was aligned to each animal’s learning point, the rate of dissociation did not differ between groups, suggesting that lOFC silencing increased the number of trials required to initiate strategy transitions without affecting their subsequent speed (**Supplementary Fig. 2b**). Consistent with this, inter- individual variability in reward-driven strategy use increased during the transition to expert reversal performance (RN→RE) in lOFC-silenced mice (**Supplementary Fig. 2c-e**). Notably, lOFC-silencing during initial learning did not affect performance (**Supplementary Fig. 2f,g**).

**Figure 2.**
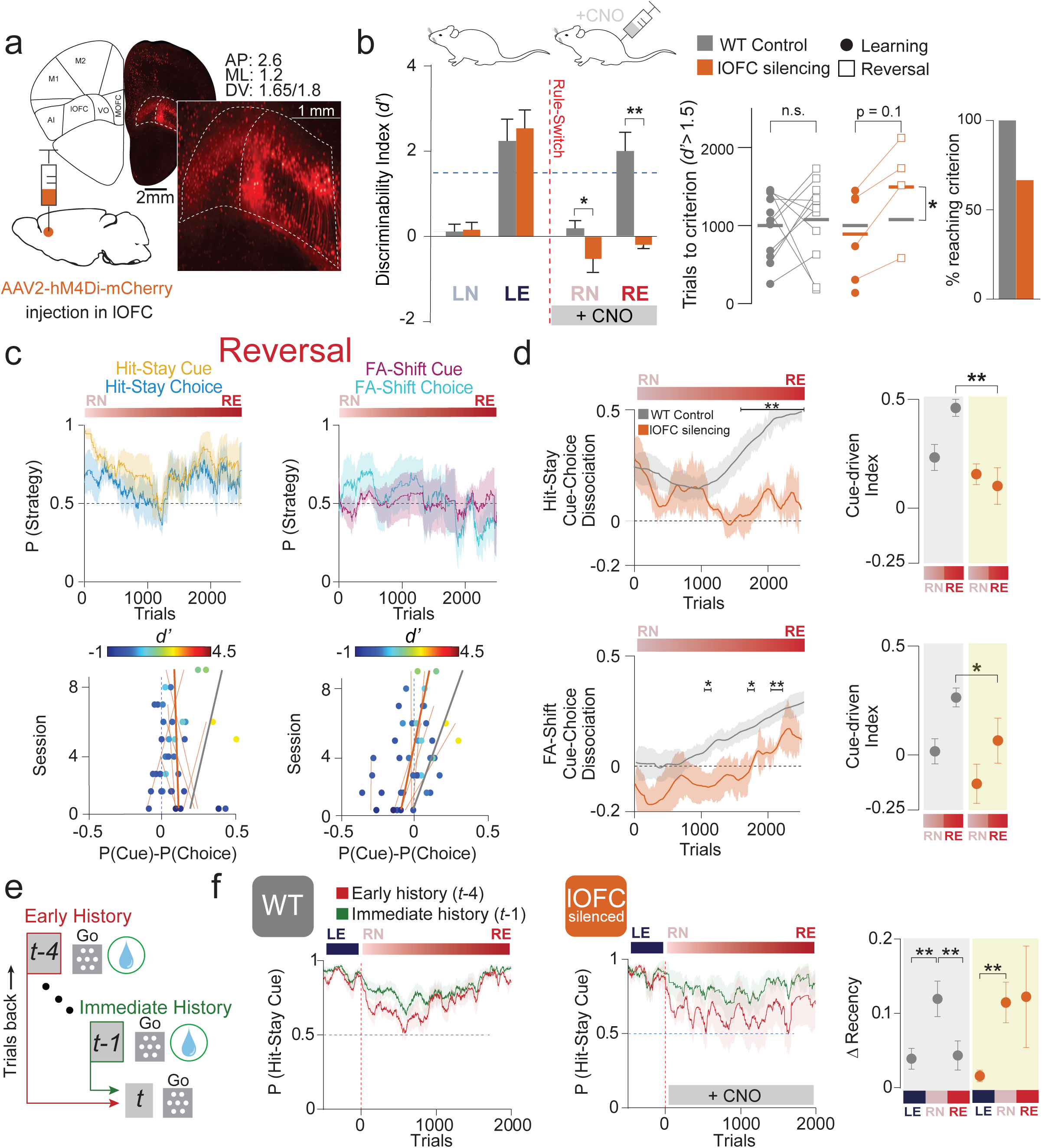
Silencing lateral OFC delays choice-to-cue strategy transitions in reward and error-based learning. **a**, Viral expression of inhibitory DREADD (hM4Di) for silencing excitatory L2/3 lOFC neurons through unilateral injection to the ipsilateral side of the barrel field. **b**, Left, comparison of behavioral performance (*d*’) of lOFC-silenced mice and WT controls aligned to the session that WT averaged performance reached the respective task phase. Neuronal silencing was achieved by daily systemic injection of CNO during reversal learning (RN→RE). Middle, comparison of trial numbers, both groups needed to reach expert criterion (*d*’ > 1.5) in initial and reversal learning. Right, bar graphs show the percentage of animals that reached expert criterion during reversal learning (*n* = 12 WT and *n* = 6 lOFC silenced mice). **c**, Top, comparison of cue and choice-based strategy dynamics from lOFC-silenced mice during reversal learning for ‘hit-stay’ and ‘FA-shift’ strategies. The curves show the mean ± S.E.M. (*n* = 4 mice). Bottom, scatterplots displaying the relation between cue-choice dissociation and session number, the color indicates d’ of the respective session. Thick lines indicate average linear fits (grey: WT, orange: lOFC silencing), and thin orange lines represent linear fits for individual lOFC-silenced mice. **d**, Dynamics and quantification of cue-choice dissociation for ‘hit-stay’ (top) and ‘FA-shift’ (bottom) strategies across task phases. **e**, Schematic depicting the definition of ‘immediate history’ (*t-1*, green) and ‘early history’ (*t-4*, red). **f**, Left and Middle, comparison of cue-dependent reward learning based on recency of experienced outcome (‘early history’ versus ‘immediate history’) in WT (left) and lOFC-silenced animals (middle) during reversal learning. Right, quantification of the difference between ‘immediate’ and ‘early’ history (ΔRecency) in salient task phases (LE, RN, and RE). Data presented as mean ± S.E.M.; *p<0.05, **p<0.01, ***p<0.001 paired two-tailed t-test, Kolmogorov–Smirnov test. Horizontal bars indicate significant differences between curves at the respective trials, cluster-based permutation test.

Thus far, analyses have focused on strategies that adjust actions following a particular action (‘choice’) or a specific stimulus (‘cue’) in the preceding trial (’immediate history’, *t*-1). However, animals can differentially weigh immediate versus distant past experiences during decision-making^25–26^, a feature previously associated with OFC function^7,27–28^. To assess the influence of longer trial histories, we analyzed trial-pairs between the current trial (*t*) and four trials back (*t*-4, ‘early-history’) as previously shown^29^ (**Fig. 2e**). Markedly, following rule-switch (RN), WT animals transiently relied more on immediate trial-history during reward-driven exploration, illustrated by a brief increase of ‘recency index’ (ΔRecency) (**Fig. 2f**, see **Methods**). In contrast, lOFC-silenced mice showed a perseveration of recency bias, marked by an absence of transition back to integrating longer trial histories during reversal learning (RN→RE) (**Fig. 2f**). Thus, intact lOFC function appears necessary for overcoming recency bias and promoting the integration of early trial experience during behavioral updating.

### Low-dimensional neural responses encode and track behavioral strategies in S1

To investigate how distinct behavioral strategies engage neural representations, we recorded neuronal population activity using in vivo two-photon Ca^2+^ imaging (**Fig. 3a**, see **Methods**). We imaged excitatory L2/3 neurons in S1, the principal brain area for somatosensory learning and integration, longitudinally across all task stages. We then applied a trial-resolution tensor component analysis (TCA) to decompose neuronal population-level representations into low-dimensional components (**Fig. 3b**)^30–31^. Each component was comprised of three factors: neuron, time, and trial. Neuron factors captured the weights of individual neurons in a component, time factors characterized the within-trial response profile, and trial factors quantified trial-by-trial gain of the response profile for a component. Based on the temporal profiles of the time factors, we identified components aligned to stimulus, action, and outcome epochs across all mice (**Supplementary Fig. 3a,b**; see **Methods**). As mice transitioned from naïve to expert (LN→LE, RN→RE), stimulus and outcome components exhibited increasingly selective trial activation (trial factor) for their corresponding task variable indicated by increased ‘component demixing’ (**Fig. 3c**, **Supplementary Fig. 3c**; see **Methods**), revealing stronger task-selectivity of these variables in S1 population-level codes (**Supplementary Fig. 3d**). Selectivity decreased immediately upon rule-switch (LE→RN) and subsequently recovered with relearning (**Supplementary Fig. 3c**). On the other hand, action-aligned components showed stable component demixing, and thus stable action-selectivity, across task phases. Collectively, these findings revealed that S1 population-level representations of stimuli and outcomes were strengthened during learning, transiently disrupted upon rule-switch and subsequently recovered during relearning, mirroring the trajectory of behavioral performance.

**Figure 3.**
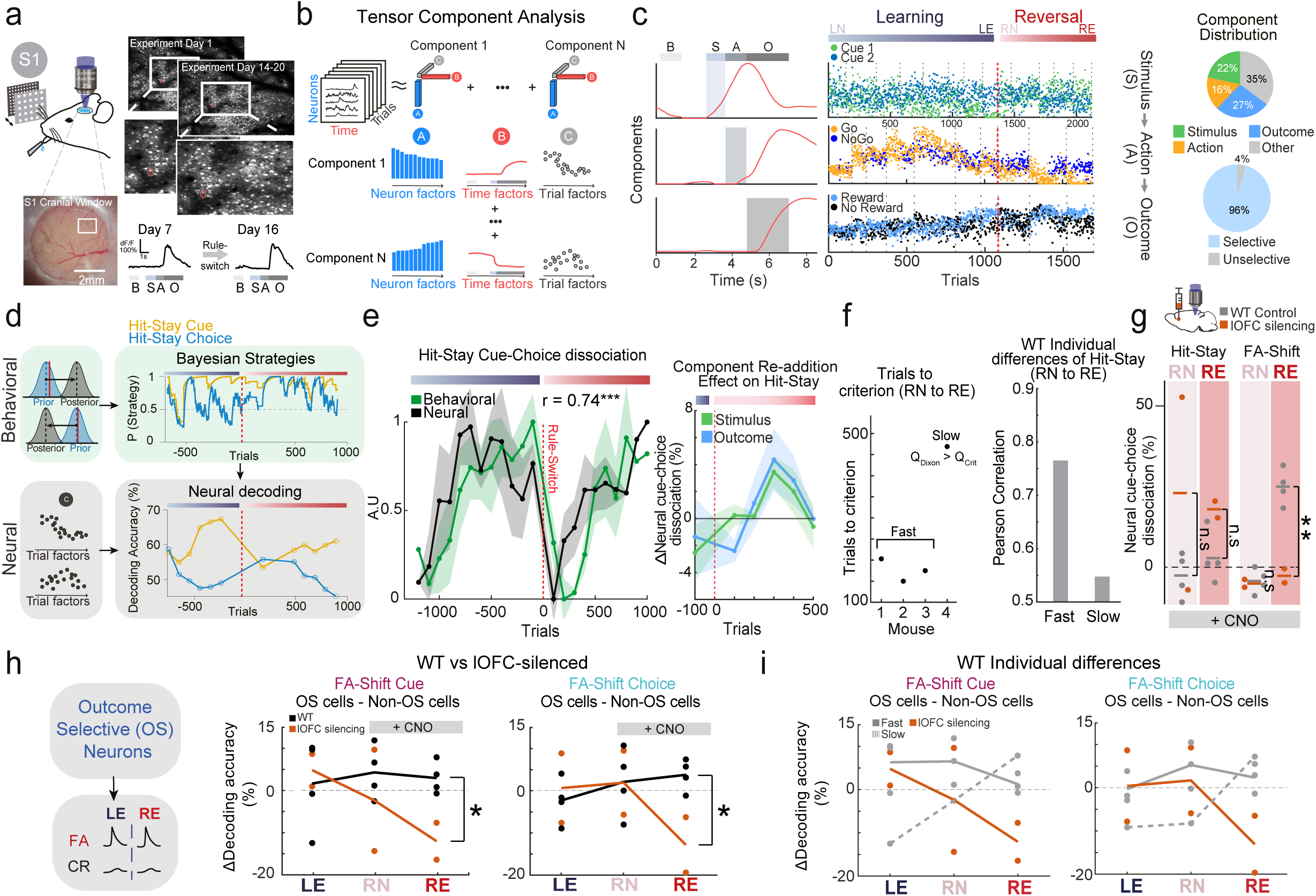
Behavioral strategy representations in S1 revealed through tensor component analysis. **a**, Left, schematic and photograph of cranial window for imaging primary somatosensory cortex (S1). Top right, 2-photon fluorescence images of excitatory L2/3 neurons imaged longitudinally over weeks of learning. Bottom right, single-trial GCaMP6f signals (ΔF/F) from one longitudinally-recorded example neuron. B, baseline; S, stimulus-presentation window; A, action window; O, outcome window. **b**, Schematic of TCA pipeline. Calcium (Ca^2+^) traces are organized into a tensor (neurons X time X trials) and decomposed into a linear sum of low-dimensional components. **c**, Left, example components of three types (stimulus, action, outcome) (2 mice). Top right, distribution of component type. Bottom right, distribution of pooled stimulus, action and outcome components by trial factor selectivity (*n* = 4 mice, 836 active neurons). **d**, Schematic of strategy-decoding pipeline (see Methods). **e**, Left, normalized behavioral cue-choice dissociation (P(‘cue’)-P(‘choice’)) and normalized neural dissociation of decoding accuracy (Accuracy(‘cue’)-Accuracy(‘choice’)) for ‘hit-stay’. Right, dissociation of decoding accuracy for ‘hit-stay’ without-minus-with stimulus or outcome components. While no single window reached significance, the second, third and fourth windows after reversal were together different from zero for both component types (p<0.05, one-tailed t-test). **f**, Left, trials to criterion (chance-performance) after reversal. A ‘slow’ adapting mouse was identified using Dixon’s Q test. Right, individual differences of behavioral and neural cue-choice dissociation correlation for ‘hit-stay’ between fast (n = 3) and slow (n = 1) adapting WT mice over reversal. r, correlation coefficient; A.U, arbitrary units. **g**, Comparison of ‘hit-stay’ and ‘FA-shift’ cue-choice dissociation of decoding performance between WT and lOFC-silenced mice after rule-switch (n = 4 WT mice, 2 silenced mice) (RN, RE). **h**, ‘FA-shift’ cue and choice decoding accuracy of outcome-selective neurons relative to non-outcome-selective neurons before (LE) and after rule-switch (RN, RE) for fast and slow adapting WT mice and lOFC-silenced mice. **i**, Same as **h** but for all WT vs lOFC-silenced mice. Data displayed as mean ± S.E.M., *p<0.05, **p<0.01, Kolmogorov-Smirnov test, one-tailed t-test, two-tailed t-test, two-tailed Wilcoxon rank-sum test, Bonferroni-Holm correction for multiple comparisons, Pearson correlation, Dixon’s Q test.

Having found population representations of stimuli, actions and outcome we asked whether behavioral strategies could be decoded directly from S1 population activity? To address this, we developed a novel strategy-decoding analysis based on TCA. We trained multinomial logistic regression classifiers to decode strategy implementation (either strategy use, avoidance, or absence) from trial factors at different task stages using Monte Carlo cross-validation (**Fig. 3d**). We found that the decoding accuracy of ‘hit-stay cue’, ‘hit-stay choice’, ‘FA-shift cue’, and ‘FA-shift choice’ strategies exceeded shuffled controls across the vast majority of trials (**Supplementary Fig. 4a,b**). Additionally, ‘hit-stay cue’ and ‘hit-stay choice’ decoding performance exceeded separately modeled strategy classification based on the prior assumption that only outcome information was leveraged in S1, demonstrating that strategy encoding could not be explained exclusively by outcome encodings (**Supplementary Fig. 4c**; see **Methods**).

To assess whether neural strategy representations tracked behavior, we compared cue-choice dissociation of neuronal decoding performance (Accuracy(‘cue’) – Accuracy(‘choice’)) with behavioral strategy use (P(‘cue’) – P(‘choice’)). We observed a strong correlation between ‘hit-stay’ cue-choice dissociation of decoding performance and strategy use (**Fig. 3e**, **Supplementary Fig. 4d**), with similar trends for ‘FA-shift’ across key task phases (LE, RN, RE) (**Supplementary Fig. 5a**), indicating that reward- and error-based strategy-related information in S1 population activity covaried with behavior. To examine the contribution of stimulus- and outcome-aligned components that covaried with performance to behavioral strategies, we recomputed decoding performance after removing these components. Excluding outcome and stimulus components reduced cue–choice dissociation, and re-introducing them restored decoding performance, particularly for ‘hit-stay’ following rule switch in the early stages of re-learning (**Fig. 3e**). This finding indicates that both S1 outcome- and stimulus-related activity may importantly provide online tracking of reward-driven strategies following a rule-switch. Notably, ‘fast-adapting’ (see **Methods**) WT mice exhibited a stronger correlation between behavioral ‘hit-stay’ cue-choice dissociation and neural decoding performance than ‘slow-adapting’ WT mice over reversal (**Fig. 3f**). This suggests ‘hit-stay’ strategy tracking by the S1 neuron population may underlie individual differences in adaptation to reversal.

Lateral OFC has been shown to play an important role in encoding decision variables relevant to behavioral strategies^32^ and shaping sensory processing^9^. To assess the causal role of lOFC feedback signals for strategy representations in S1, we compared the cue-choice decoding performance dissociation from excitatory (L)2/3 neurons in WT and lOFC-silenced mice for ‘hit-stay’ and ‘FA-shift’ strategies. During reversal learning, WT mice exhibited a significant increase in ‘FA-shift’ decoding dissociation, which was absent in lOFC-silenced mice (RN→RE) (**Fig. 3g**). In contrast, ‘hit-stay’ decoding dissociation did not differ between these groups (**Fig. 3g**). This underscores the general importance of functional lOFC signaling for strategy representations in S1 populations, but specifically for error-related strategy representations during relearning. We previously identified an outcome-selective subpopulation of excitatory neurons in S1 that were specifically modulated by lOFC feedback signal^9^. We examined the specific role of these neurons in population-level ‘FA-shift’ strategy representations (**Supplementary Fig. 5b,c**). In WT animals, outcome-selective neurons, compared to a random sample of non-outcome-selective control neurons of the same number, contributed more to ‘FA-shift’ cue and choice strategy decoding in the reversal expert phase (RE) for WT mice (**Fig. 3h**). Markedly, decoding performance of outcome-selective neurons in ‘slow’ adapting WT mice showed poorer, though non-significant, encoding in RN, but not RE (**Fig. 3i**), suggesting a delayed remapping of these neurons’ strategy representations. Further analysis revealed that decoding performance was significantly reduced for outcome-selective neuronal responses in lOFC-silenced mice compared to WT outcome-selective neurons (**Fig. 3h**) but not compared to WT stimulus-selective neurons (**Supplementary Fig. 5d**). These results suggest that lOFC feedback signals guide error-guided strategy representations by specifically modulating outcome-selective excitatory neuronal subpopulations in S1 during reversal learning.

### Outcome-selective S1 neurons integrate strategy-relevant outcomes

How exactly do outcome-selective neurons in S1 contribute to adaptive strategies during behavior? One possibility is that these neurons may integrate reward- and error-history with current experience to support strategy representations during learning.

To examine strategy tracking mechanisms in outcome-selective S1 neurons, we calculated each neuron’s selectivity for respective strategy-specific outcomes in trial *t*. These outcomes indicated whether an animal used or avoided a given ‘cue’ strategy (‘hit’ vs ‘miss’ for ‘hit-stay cue’ (‘hit-selective’); ‘FA’ vs ‘CR’ for ‘FA-shift cue’ (‘FA-selective’)). Based on their response profile, outcome-selective S1 neurons were sorted (see **Methods**) into three candidate strategy neuron subpopulations (‘hit-selective’, ‘FA-selective’, ‘mixed-selective’) (**Fig. 4a, Supplementary Fig. 6a**). We calculated reward- and error-history modulation indices (RHMI and EHMI, respectively) to quantify outcome-history-dependent modulation of single-neuron responses for each of these three subpopulations. For example, two consecutive reward outcomes indicate use of ‘hit-stay cue’ reflected in a RHMI deflection (>0 enhanced response, and <0 suppressed response; similar deflection for EHMI for the ‘FA-shift cue’ strategy) (**Fig. 4b**). ‘Hit-selective’ neurons in WT mice showed a stable increase in RHMI, effectively switching to being positively modulated by prior rewards, demonstrating their selective responses to the use of ‘hit-stay cue’ immediately after rule-switch (RN, RE) (**Fig. 4b**). ‘FA-selective’ neurons, on the other hand, showed an increase in EHMI as they selectively responded to the use of ‘FA-shift cue’ only in the late reversal phase (RE) (**Fig. 4c**), reflecting mice’s slower behavioral cue-choice dissociation for error- than reward-driven learning. Remarkably, the history modulation of both neuron subpopulations changed sign following lOFC inactivation (**Fig. 4b,c**), resulting in a switch from positive to negative modulation of ‘hit-selective’ neurons in RN, and of ‘FA-selective’ neurons in RE. Importantly, both the ‘hit-selective’ and the ‘FA-selective’ subpopulations displayed high specificity for their respective strategies (‘hit-selective’ for ‘hit-stay cue’; ‘FA-selective’ for ‘FA-shift cue’). Neither subpopulation showed modulation by outcome-history relevant to the alternate strategy, as evidenced by the absence of cross-modulation (**Fig. 4d,e**). Positive history modulation from prior rewards or errors was not observed for non-selective neurons in any task phase (**Supplementary Fig. 6b,c**), further highlighting the functional specificity of outcome-selective subpopulations. The ‘mixed-selective’ cells showed a transient increase in RHMI during early reversal (RN), but a transient decrease in EHMI (RN) and subsequently increased, though non-significantly, EHMI in the late reversal phase (RE) (**Supplementary Fig. 6d,e**), broadly indicating a transition from reward tracking in RN to error tracking in RE, mirroring mice’s slower behavioral cue-choice dissociation for error- than reward-driven learning.

**Figure 4.**
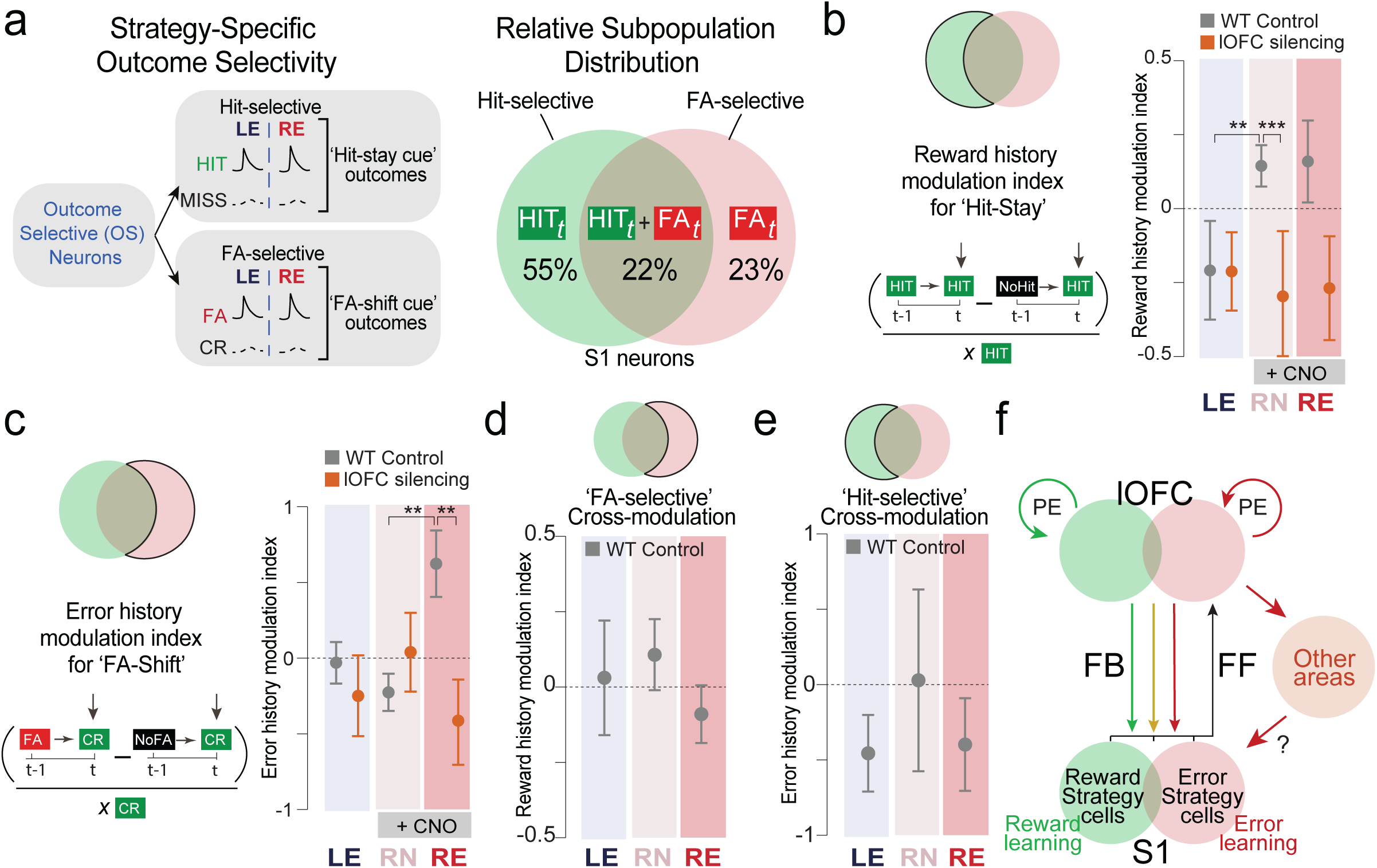
Lateral OFC-dependent integration of strategy-specific past and current outcome variables. **a**, Left, defining functional subpopulations of neurons showing specific strategy-relevant outcome-selectivity in trial *t* (’hit-selective’, ‘hit’ versus ‘miss’; ‘FA-selective’, ‘FA’ versus ‘CR’). Right, relative distribution of ‘hit-selective’, ‘FA-selective’ and ‘mixed-selective’ neuron subpopulations. **b**, Left, calculation of the ‘hit-stay cue’ associated reward history modulation index (RHMI). Neural responses of ‘hit’ trials (defining strategy use in trial *t*), preceded by a ‘hit’ trial, and those preceded by a ‘non-hit’ trial, are subtracted and divided by the average ‘hit’ trial response to indicate reward-driven history modulation (positive values) for ‘hit-stay cue’ strategy-specific variables. Right, comparison of RHMI for ‘hit-selective’ neurons in WT and lOFC-silenced animals across task phases (LE, RN, and RE). **c**, Left, calculation of the ‘FA-shift cue’ associated error history modulation index (EHMI). Neural responses of ‘CR’ trials (defining strategy use in trial t) preceded by a ‘FA’ trial and those preceded by a ‘non-FA’ trial are subtracted and divided by the average ‘CR’ trial response to indicate error-driven history modulation (positive values) for ‘FA-shift’ strategy-specific variables. Right, comparison of EHMI for ‘FA-selective’ neurons in WT and lOFC-silenced animals across task phases (LE, RN, and RE). **d**, RHMI for ‘FA-selective’ neurons (cross-modulation) in WT animals across salient task phases (LE, RN, and RE). **e**, EHMI for ‘hit-selective’ neurons (cross-modulation) in WT animals across salient task phases (LE, RN, and RE). **f**, Schematic showing potentially differential lOFC feedback (FB) targeting distinct subpopulations of S1 neurons, embedding the ability to specifically track strategy-relevant variables. Feedback signals to S1 from other integrative cortical or subcortical areas may also contribute. FF, feedforward; PE, prediction error. *n* = 3 WT mice and *n* = 2 lOFC-silenced mice for all plots. Data presented as mean ± S.E.M.; *p<0.05, **p<0.01, ***p<0.001, two-tailed t-test.

Taken together, these results reveal distinct outcome-selective S1 subpopulations whose activity exhibits strategy-specific history modulation. Lateral OFC differentially attenuated these reward- and error-history signals in these subpopulations, highlighting the multiplexed nature of the frontal feedback teaching signals implementing behavioral flexibility (**Fig. 4e**).

## Discussion

Unraveling how brain-wide interactions support animal exploration during adaptive behavior remains a core challenge in neuroscience. In recent years, we have seen a renewed focus on understanding animal behavior using quantitative approaches^33,34^. When integrated with advanced large-scale, high-resolution neural recordings^35–36^ and causal manipulations^37–38^, these approaches provide a powerful framework for dissecting neural mechanisms underlying behavioral diversity. The lOFC has been identified as a key brain area contributing to critical variables associated with behavioral strategies. OFC neurons construct meaningful associations between stimuli, actions and outcomes^24,39^, track trial history^28,40^, and update belief states in inference-based tasks^41–42^. In our study, we built on these observations and revealed a novel role of lOFC-mediated cognitive control whereby, via the broadcasting of control signals to sensory areas, lOFC guides animals’ sensory strategies during flexible behavior. Mice trained on a tactile reversal learning task switched from action-to stimulus-driven strategies during learning and reused learnt strategies during rule reversal. Chemogenetic silencing of the lOFC impaired both reward- and error-driven learning strategies to varying degrees. We identified distinct subpopulations of reward-and error-driven strategy neurons in the superficial L2/3 of S1, and modulation of these subpopulations by past outcome history during relearning depended on lOFC. This highlights the distributional nature of the lOFC feedback signal that engages S1 neurons, thereby impacting reward- and error-guided strategies. These findings bridge behavioral strategy inference with circuit-level interactions between the frontal and sensory cortex.

### Action-driven to stimulus-driven strategy exploration during tactile reversal learning

Mice adapt their behavior by closely sampling sensory stimuli, prior choices and associated rewarding and aversive outcomes, yet how they use this observable and latent information to adjust their behavioral strategies during adaptive learning remains a key open question. We implemented a trial-resolution Bayesian strategy analysis to assess how animals use specific strategies during learning. Unlike classical psychophysical performance measures like sensitivity or discriminability index, our strategy analysis provides a richer and quantitative insight into behavior, revealing how individual animals explore action/stimulus-driven strategies in the face of changing contingencies. Leveraging trial-resolution analysis is therefore critical given the non-stationary nature of choice behavior in changing environments^17,19^. We found that mice transitioned from using action- to stimulus-driven strategies during learning and relearning, providing evidence for attention-theory frameworks that argue animals gradually direct attention towards reward-predictive task dimensions over learning^43^. The use of action-driven strategies in the early stages of initial learning was expected, given that licks were rewarded during pre-training and the animals’ motivational state.

Due to the initial lick bias and the collective transition to stimulus-based exploration, mice’s use of strategies covaried through initial learning (LN, LE) but decorrelated following the rule-switch (RN). While most animals coherently relied more on action-based strategies immediately after rule-switch, they varied considerably in their retention of stimulus-driven exploration. This suggests that following contingency reversal in RN, attention predominantly returns to the attended feature (action) during the LN task phase, but is accompanied by increased attentional variability. Interestingly, this is also the only task phase (RN) in which a subset of animals showed greater dissociation between error-driven and reward-driven learning. This further highlights how an abrupt change in contingency results in a transient, broader, and more diverse strategy use, before shared experience in the same contingency drives reconvergence across animals. This result holds key significance for Bayesian learning frameworks, suggesting that adjustments in learning rate, inferred context and belief due to a change in contingency may co-occur with an increase in individual variability of these cognitive variables, before they re-converge on the new task structure^44–46^. Such adaptive behaviors require the involvement of prefrontal areas of the neocortex, and especially the OFC. We found that pharmacogenetic silencing of the lOFC during re-learning impaired the transition from action- to stimulus-driven choices (RN→RE), initially through reward-guided learning strategies in RN and later through error-guided strategies in RE. Saliently, reward- but not error-driven strategies exhibited higher individual variability in RE for lOFC-silenced mice. This suggests lOFC stabilizes reward more than error-based learning across individuals in RE. While studies have reported a reward-error dissociation in relearning following lOFC silencing^47^, the performance-phase specificity we revealed has not been reported before to our knowledge. These results highlight the dynamic importance of lOFC in specific performance phases for reward- and error-based learning, strategy stability across individuals, and the transition from action- to stimulus-driven choice.

### Decoding of reward- and error-based strategies in primary sensory cortices

The S1 is classically considered to predominantly encode features of tactile stimuli^48–50^. We implemented a trial-resolution dimensionality-reduction approach, tensor component analysis (TCA), to disentangle overlapping stimulus-, action-, and outcome-related signals and track their evolution across learning^30–31^. Additionally, this approach allowed us to examine whether contributions from sensory and non-sensory task variables were present at single-trial resolution in the activity of hundreds of longitudinally recorded L2/3 neurons in S1. By aligning trial-resolved TCA with multinomial logistic regression decoding, we directly linked low-dimensional neural population structure to latent behavioral strategies on a trial-by-trial basis, bridging dimensionality reduction with strategy inference, extending prior work on dimensionality reduction^30,51^, and behavioral strategy inference^17,19^ to a unified neural–behavioral framework. This powerful analytical approach revealed representations not only of tactile stimuli but also of actions and outcomes^14,52–53^. The selectivity of trial-factor activations for stimuli and outcomes, but not actions, covaried with behavioral performance, indicating that selective stimulus and outcome responses in S1 may support stimulus-outcome learning. In particular, we found that the observed TCA components (stimulus, action, and outcome) of S1 responses represented information necessary to track the use of reward- and error-based strategies. We trained a linear classifier on TCA trial factors to decode whether S1 population responses encoded strategies and how these might evolve during adaptive learning. We found that selective S1 neurons continuously represented and tracked reward- and, more broadly, error-driven strategies during task learning. Importantly, fast-adapting mice demonstrated stronger, though non-significant, reward-driven strategy tracking in S1 neurons than slow mice over relearning, linking the quality of S1 strategy tracking to behavioral performance. To our knowledge, the existence of strategy representations in S1, dependent on the integration of multiple variables across trials, has not been reported previously. These non-sensory codes in S1 add to the growing evidence for diverse cognitive codes and computations in sensory cortices^3^. These range from complex sensory processes, such as texture discrimination^54^, to abstract higher-order codes of reward^13^, decision variables^14^, trial-history^9^, timing, surprise and prediction errors^15,55^. These patterns might similarly engage other primary sensory cortices, such as the encoding of stimulus reward associations and prediction errors in V1^56–57^ and A1^58–59^. Together, our findings highlight the complexity of rodent S1 and uncover its dynamic tracking and representation of behavioral strategies during adaptive learning.

### Fractionating lOFC-dependent behavioral strategy representations in S1

Our previous work revealed that a small subpopulation of outcome-selective L2/3 excitatory neurons in S1 remapped response selectivity during reversal learning via lOFC teaching signals^9^. These results, together with our finding that lOFC-silencing affected strategy use during relearning, prompted our inquiry into whether lOFC signals could also instruct S1 strategy codes. We found that pharmacogenetically silencing lOFC disrupted stimulus-driven error-, but not reward-strategy codes in the S1 neuronal population during RE. This was surprising as we hypothesized that both error- and reward-strategy codes would depend on intact lOFC signaling, due to the observed behavioral effect of lOFC silencing on both processes^60–62^. Additionally, encoding of behavioral variables relevant for both strategies^61,63^ as well as unsigned prediction error signaling^64^ were previously reported in lOFC, which could support such teaching signals. Our TCA analysis revealed an lOFC-dependent S1 subpopulation (‘FA-selective’) that showed stronger error-driven strategy encoding than ‘non-outcome’ selective or ‘stimulus-selective’ neurons in RE. Strikingly, lOFC-silencing reduced ‘FA-selective’ neurons’ encoding of error-guided strategy to well below ‘non-outcome selective’ neurons in RE, suggesting lOFC plays a crucial role in the optimal functioning of these neurons after reversal. Together, behavioral effects of lOFC-silencing were evident for both reward- and error-guided strategies across phases, whereas the S1 neuronal decoding reduction was most pronounced for stimulus- driven error strategies during late reversal, and S1 strategy codes appear to be heterogeneously distributed within the L2/3 population. This dissociation reinforces the existence of specialized yet partially overlapping neural circuits in S1. Importantly, the lOFC not only projects to S1^9^ but also to several other primary sensory cortices in mice, with lOFC teaching signals instructing A1^59^, olfactory cortex^66^, gustatory cortex (Unpublished observation, Erin Rich lab), and the V1^67^. It is possible that positive, negative, and multiplexed valence-encoding lOFC neurons^68–70^ fall into relatively distinct functional clusters, which, in turn, modulate strategy-specific response selectivity and regulate sensory strategies. Importantly, strategy representations could also be observed during the initial learning phase (LN→LE), during which lOFC silencing showed no effect on behavioral performance or strategy use (see also **Supplementary Fig. 2**). This indicates that lOFC input is necessary to maintain error-driven strategy representations only after sudden changes in contingency, in line with lOFC’s proposed involvement in signaling a contextual switch. Taken together, lOFC-dependent embedding of strategy-tracking or other associative codes could therefore be a generalizable mechanism across primary sensory cortices, opening a fascinating line of enquiry that future studies could address.

### Outcome-selective S1 neurons integrate strategy-specific variables to support flexible behavior

What cellular mechanisms could support strategy representations in S1? We reasoned that (1) contribution to population representations of strategies, (2) the ability to track and integrate task-history information across trials, and (3) a disruption of these functions following silencing of lOFC activity during task-switching, are key criteria. We identified S1 ‘outcome-selective’ neurons that satisfied all these criteria and investigated the cellular mechanisms underlying their strategy codes. We identified specialized ‘hit-selective’ and ‘FA-selective’ neuronal subpopulations that selectively responded to strategy-specific outcomes following the use or avoidance of a ‘cue’ strategy. Furthermore, the history modulation of these subpopulations by strategy-specific outcomes was observed during relearning and depended on lOFC input. Because history modulation effects appeared during relearning, not LE, this suggests modulation occurs specifically due to prior strategy outcomes rather than increased arousal or motivation from prior licking. Importantly, we found that the outcome-selective neurons drive both population-level decoding of strategy and single-cell-level integration of outcome-history. This provides a mechanistic link between top-down orbitofrontal signaling and a population-level representation of strategy use via a single-neuron-based mechanism in specialized subpopulations in S1. This finding may have significant implications for reconciling the single-neuron doctrine and the competing population doctrine^71^, revealing how population representations of strategies are supported by single-neuron mechanisms within strategy-specific subpopulations. An interesting emerging question to address would be whether these subpopulations in S1 are driven by vectorized subpopulation-specific top-down signals from lOFC, or rather one homogenous, global signal that is vectorized within S1 involving local microcircuitry. We previously reported similar hierarchically organized circuits in the lOFC involving VIP+ disinhibitory interneurons^65,72^; sensory cortices may mirror similar circuit architecture^67^ to accommodate flexible learning.

Finding cellular mechanisms for strategy encoding in S1 was unexpected, as it requires neurons in these subpopulations to encode multiple task variables and integrate them across temporal scales during learning – features that are traditionally assigned to prefrontal neurons. This is a rather unique ability of outcome-selective neurons in S1, to our knowledge, and it provides a new mechanistic insight into the cellular-level codes of reward- and error-driven learning in the sensory cortex. While the exact function of such representations in a primary sensory area remains unclear, the strong contribution of outcome-selective neurons to both population representation and single-cell level mechanisms indicates a more retrospective “tracking” of the strategy used in that trial when the outcome is received, evocative of recently proposed retrospective cognitive maps^73^. These coding schemes also resonate with computational theories of reinforcement learning, in which both prediction errors and accumulated reward histories are essential for updating internal models of the environment. Importantly, our findings demonstrate that such computations are not restricted to the prefrontal cortex but are distributed across cortical circuits, with S1 playing an unexpected role.

Our study suggests that pre-existing non-sensory codes, such as actions, outcomes, and strategy-related information, in S1 can be further attuned by top-down instructive signals depending on the behavioral context. This S1 information may be relayed via a bottom-up signal, with potentially different sources and targets depending on the current state of the environment or the animals’ knowledge of it. Taken together, our findings reveal that complex exploratory learning strategies guide animals on flexible learning tasks. Such strategies can be dynamically decoded from neuronal responses in S1 and distinctly modulated by top-down lOFC signals, at both the mechanistic cellular and population levels. These findings further suggest a revision of the classical hierarchical bottom-up model of sensory processing and suggest an interactive model that incorporates both bottom-up and top-down neural interactions^3,74^, with dynamically changing partners based on the animal’s task state or the environment. Our results suggest S1 might act as a dynamic reservoir where orbitofrontal feedback imprints strategy- and outcome-related representations. We hypothesize these S1 representations provide a rich substrate for downstream circuits to read out to guide long-term strategy adaptations, support internal refinements of bottom-up sensory signaling, and facilitate the transformation of goal-directed to habitual behavior.

## Acknowledgements

We thank the past and current members of the Adaptive Decisions Lab for their critical comments and discussions on this manuscript. We thank Professor Fritjof Helmchen for his support.

This work was supported by a Wellcome Trust Career Development Award (227782/Z/23/Z), a Wellcome Trust institutional strategic award at Newcastle University, a Royal Society research grant (RGS\R2\202155), and, previously, a Novo Nordisk Foundation Young Investigator Award (all to A.B.) and a Reece Foundation PhD Fellowship (to R.R. and A.B.). A.B. is affiliated with the Digital Environment Research Institute at QMUL and the Department of Psychiatry at the University of Oxford. The authors declare no conflict of interest.

## Author contributions

A.B. conceived and designed the study. J.T. carried out all experiments. R.R. developed novel data analysis pipelines. S.M. and M.D.H. developed the Bayesian strategy analysis pipeline. J.T. and R.R. analyzed data. J.T., R.R., and A.B. wrote the manuscript with comments from all authors.

