## Supplementary Material for "Distinct Orbitofrontal Feedback Signals Shape Sensory Behavioral Strategies during Flexible Learning"

### Methods

#### Animals

All experimental procedures were conducted in accordance with the requirements of the United Kingdom Animals (Scientific Procedures) Act 1986, and the Federal Veterinary Office of Switzerland, and were approved by Newcastle University's Animal Welfare and Ethical Review Board (AWERB) and the Cantonal Veterinary Office in Zurich. Experiments were conducted under the authority of Home Office Project License PABAD450E and license numbers 285/2014 and 234/2018. Mice were housed at 24°C in a 12-h reverse dark-light cycle (07:00 to 19:00). At the end of the experiment, the mice were deeply anaesthetized, transcardially perfused and euthanized by exposure to $CO_2$ in their home cage. For behavioral experiments, we used wild-type (WT) C57BL6/J mice (n = 20 mice). For imaging neurons in S1 (n = 6 mice), we used Rasgrf2-2A-dCre:CamK2a-tTA:TITL-GCaMP6f triple transgenic mice, which express GCaMP6f in excitatory neocortical L2/3 neurons. To generate triple transgenic mice amenable to two-photon imaging, double transgenic mice carrying CamK2a-tTA (Jackson Laboratories no. 016198[32]) and TITL-GCaMP6f (Jackson Laboratories no. 024103[33]) were crossed with a Rasgrf2-2A-dCre line (Jackson Laboratories no. 022864[34]). The destabilized Cre recombinase expressed under the control of the Rasgrf2-2A promoter was stabilized by trimethoprim (TMP, Sigma T7883) to render it functional. TMP was reconstituted in dimethyl sulfoxide (DMSO, Sigma 34869, 100 mg/ml), freshly prepared before each induction, and administered 2 weeks before surgery. During induction, mice were given a single intraperitoneal injection (150 mg TMP per g body weight diluted in 0.9% saline solution) using a 29-G needle. Additionally, in a few mice, we injected AAV viral vectors into cortical L2/3 expressing GCaMP6f under the hSyn promoter. For causal pharmacogenetic manipulations, both wild-type (WT) and L2/3-GCaMP6f mice were used. All efforts were made to minimize suffering.

**Water scheduling**

On the day before behavioral training started, mice were put on a water scheduling regimen, on which they remained for the duration of the experiment. A baseline weight was calculated based on the average weight of the prior three days. Animals were weighed every day before training and kept at 85-90% of the calculated baseline weight. During training, the amount of reward received was recorded. If insufficient water was provided during the training (40 ml/kg/day) animals were allowed access to water after training. During weekends, animals were always provided with ad libitum access to water.

**Reversal learning task**

Mice were extensively handled and familiarized with both the experimenter and the experimental setup. On the day before behavioral training started, mice were put on a water-scheduling regimen, on which they remained for the duration of the experiment. During the experiment, in each trial, an auditory cue (two beeps at 2 kHz, 100 ms duration with a 50 ms interval) signaled the beginning of each trial. After the pre-stimulus window (3 s), one of two possible sandpaper textures (P100 as a rough 'Go' - texture or P1200 as a smooth 'No-Go' - texture) was presented to the whiskers for the duration of 1 s. After the withdrawal of the texture, an additional auditory tone (response cue; 4 beeps at 4 kHz, 50 ms duration with a 25 ms interval) signaled the start of the 'response window' (1 s), during which mice could indicate their choice by either licking or withholding a response. Licking for the 'Go' texture ('hit') was rewarded with sucrose water, while licking for the 'No-Go' texture ('false alarm', FA) was punished with a brief exposure to white noise. Reward and punishment were both omitted after withholding a response to either the 'Go' ('miss') or 'No-Go' texture ('correct rejection', CR)[9,75]. Performance was measured for each session using the discriminability index, *d'*, which is calculated by $Z(N_{hit}/(N_{hit}+N_{miss})) - Z(N_{FA}/(N_{FA}+N_{CR}))$ where Z is the probit function and $N_x$ is the number of trials with an outcome *x*. After mice showed an improvement in performance from naïve to expert during the initial learning phase (LN to LE) and displayed stable expert performance (reaching $d' \geq 1.5$ for three consecutive sessions), the stimulus-outcome associations were reversed. Thus, mice needed to adapt their behavior and learn to respond to the new ‘Go’ texture (P1200) while withholding licking for the new ‘No-Go’ texture (P100) to recover their performance from the reversal naïve to reversal expert stage (RN to RE). For naïve sessions (LN, RN), the first session with a general engagement of > 50% (usually first or second session) was chosen. For the expert phase the last session with *d'* > 1.5 was chosen (200–300 trials/session/day).

**Bayesian strategy analysis**

To examine how mice used behavioral strategies trial-by-trial, we analyzed choices with a Bayesian evidence accumulation model[17]. This analysis estimated the probability that a behavioral strategy is being used on any given trial, given the history of choices up to that trial. In short, the analysis estimates the posterior probability distribution, *P*, that a strategy *j* is being used on a trial *t* given the choices in all history from trial 1 to *t*. By leveraging Bayes theorem and choosing a uniform prior distribution, one can derive $P(\text{strategy}_j(t)|\text{choices}(1{:}t))$ which is the posterior distribution at every trial *t*. The posterior probability is updated based on the *likelihood* - how in line the choices made from trial 1 to *t* are with the strategy *j* - and the *prior* – the original probability estimate for using that strategy (which is a uniform distribution at trial 0). After the first trial, the posterior of trial *t* can be calculated using the posterior of *t*-1 as the *prior,* for all subsequent trial updates. To extract a probability estimate that a given strategy, $\mathrm{P}(\text{strategy}_j, t)$, is being used, we extract the probability that maximizes the Posterior probability distribution function.

$$P(\mathrm{P}(\text{strategy}_j, t)) = Max(\, P\left(\text{strategy}_j(t) \middle| \text{choices}(1{:}\, t)\right)\,)$$

We defined four key strategies, ‘hit-stay cue’, ‘hit-stay choice’, ‘FA-shift cue’ and ‘FA-shift choice’. Detailed definitions of these strategies are as follows: 1) If the previous trial was a ‘hit’ and has the same stimulus as the current trial, a ‘hit-stay cue’ strategy is used if in the current trial the same choice is used as in the previous trial, and ‘hit-stay cue’ is avoided if a different choice from the previous trial is used. 2) If the previous trial was a ‘hit’, a ‘hit-stay choice’ strategy is used if in the current trial the same choice is used as in the previous trial. 3) If the previous trial was a ‘FA’ and has the same stimulus as the current trial, a ‘FA-shift cue’ strategy is used if a different choice is used in the current trial than was used in the previous trial. 4) If the previous trial was a ‘FA’, a ‘FA-shift choice’ strategy is used if the current trial uses a different choice than the previous trial.

**Cue-choice dissociation**

To measure if an animal's behavior was more 'choice-driven' (current choice based on previous choice association with outcome), we calculated P(‘hit-stay choice’) and P(‘FA-shift choice’) for each mouse over the whole task. To measure 'cue-driven' behavior (current choice based on previous stimulus association with outcome), we calculated the level of 'cue-choice dissociation' over all trials for ‘hit-stay’ and ‘FA-shift’ for each individual mouse by subtracting the ‘choice’ from the ‘cue’ strategy probability at each trial. Dissociation curves from individual animals were then averaged over animals. To quantify how 'cue-driven' an animal's behavior was in a specific task phase (LN, LE, RE, and RE), we calculated the average 'cue-choice dissociation' for that respective session.

**Simulations of strategy execution**

To determine how specific dynamics of ‘hit-stay’ and ‘FA-shift’, cue and choice strategies reflect stimulus- or action-based decisions, we created artificial agents with three predefined choice behaviors: ‘action-driven’, ‘stimulus-driven’ and ‘mixed’. During the 'action-driven' phase, agents used ‘hit-stay choice’ and ‘FA-shift choice’ when possible, but otherwise a random choice was executed (e.g. a ‘hit’ would always be followed by a ‘hit’ or ‘FA’ depending on the stimulus presented in the second trial, while the decision after a correct rejection or miss would be random). During the ’mixed’ phase, the action resulting in a strategy defining outcome (‘hit’ for ‘hit-stay’ and ‘FA’ for ‘FA-shift’) is only repeated if the same stimulus was presented in trial *t* and *t*-1, otherwise the agent chooses a random action (e.g. for 'hit-stay' a 'hit' is always followed by another 'hit' if the same stimulus is presented again, and by a random outcome if the other stimulus is presented (false alarm or correct rejection). In trials following a strategy-irrelevant outcome, the action is chosen randomly (e.g. for 'hit-stay', the decision after a false alarm, correct rejection or miss would be random). In the 'stimulus-driven' phase, the agent chooses the correct action for the presented stimulus in trial *t*, independent of information from the previous trial, thus representing a post-exploration expert level. Each phase consisted of 500 trials, and the three phases were simulated in succession. In an additional simulation, we integrated a gradually decreasing, initially highly probable 'lick bias' (90%), overwriting the predefined choice structure and forcing the agent to respond with a 'lick'. Additionally, simulations considered a 'disengagement probability' (10%), overwriting the predefined choice structure and forcing the agent to respond with a 'no lick'. All simulations were run 6 times (6 agents with lick bias, 6 agents without lick bias), strategy probabilities were extracted from all agents individually as described above for the mice.

**Trials to criterion**

Trials to *d'* > 1.5 was used as the criterion to characterize the number of trials required to reach expert performance during initial learning and reversal learning. Trials to chance performance *d'* = 0 after reversal was used to characterize the number of trials required to stop perseverating with the pre-reversal contingency.

**Experience-recency index**

To evaluate the influence of immediate (*t-1*) compared to early experience, we used trial pairs of the current trial (*t*) and four trials back (*t-4*, 'early history'). To quantitively measure the influence of different history use in each task phase, we calculated an 'experience-recency index' ('ΔRecency') by calculating the difference of strategy probabilities between immediate and early history for the LN, LE, RN and RE phases:

$$\Delta \text{Recency} = mean(\text{P}_{t-1}(\text{strategy}, \text{TP})) - mean(\text{P}_{t-4}(\text{strategy}, \text{TP}))$$

Where $P_{t-n}$ is the probability of a strategy using an outcome at trial $t - n$ to make a choice at trial $t$, TP are the task phase trials - (LN, LE, RN, RE). A positive value indicates a higher relevance of the 'immediate' history for the animals' decision, a negative value indicates a higher relevance of 'early' history, and a value around zero indicates no influence of recency.

**Virus injection**

Mice were briefly anesthetized with isoflurane (2% in oxygen) in an anesthesia chamber and transferred to a stereotactic frame (Kopf Instruments), the cranium was secured with ear bars, and eyes were covered with vitamin A cream (Bausch & Lomb). During the surgery, anesthesia was maintained with 0.8–1.2% isoflurane and the body temperature was maintained at ~37 °C and the eyes were protected with ointment. After opening the skin and cleaning the cranial surface, the lOFC location was identified based on stereotactic coordinates, 2.6 mm anterior and 1.2 mm lateral from bregma[76]. A small hole was drilled, and two injections (1.65 mm and 1.85 mm from dura mater) were made using a 10 ml a glass capillary. In each injection site, 70 nl of the virus was injected at a rate of 50 nl/min at each site, and kept in place for an additional 5 minutes to minimize backflow. A stainless-steel head post was fixed to the skull with dental cement, and the skin was reattached to the implant.

**Cranial window**

Mice were implanted with a cranial window under controlled anesthesia (as described above)[75,77]. A 4 mm craniotomy was performed, and a 4 mm round cover glass was placed on top of the dura mater. The edges were sealed with ultraviolet-light-curable dental acrylic cement (Ivoclar Vivadent). A stainless-steel head post was fixed to the skull with dental cement, and skin was reattached to the implant. Mice had at least one week of recovery before pre-training handling started.

**Drugs**

For chemogenetic inhibition experiments, inhibitory DREADDs (hM4Di), CaMKIIα-hM4D(Gi)-mCherry, were expressed in excitatory L2/3 lOFC neurons through virally mediated transfection (see Virus Injection). hM4Di was activated via intraperitoneal (i.p.) injection of clozapine N-oxide (CNO dihydrochloride, 1–5 mg/kg, Tocris Bioscience, Catalog number 4936) 25-30 minutes before each training session.

**Intrinsic signal optical imaging**

The recording site, barrel cortex in the primary somatosensory cortex (S1), was identified using intrinsic signal optical imaging under light anesthesia (0.8-1%). Whisker stimulation evoked signals were imaged at 630 nm and collected with a CMOS Python camera (The Imaging Source DMK 23UP1300; 8/10 bit; 1,280×1,024 pixels, 4.8μm pixel size, 10-Hz frame rate). Stimulus-responsive S1 cortical areas were detected by measuring differences between repeated post-stimulation responses (15 trials) and an unstimulated baseline (50 frames; expressed as $\Delta R/R$). For recording site selection, reference surface vasculature images were obtained using a 546 nm LED.

**Two-photon imaging**

Neurons were imaged using a two-photon microscope from Neurolabware equipped with a Ti: Sapphire laser system (approximately 100 femtosecond laser pulses; Mai Tai HP, Newport Spectra Physics), a water-immersion 16X Olympus objective (340LUMPlanFI/IR, 0.8 numerical aperture, NA), galvanometric scan mirrors (model 6210; Cambridge Technology) and a Pockels Cell (Conoptics) for laser intensity modulation. Virally expressed GCaMP6f in L2/3 neurons in S1 were imaged at 940 nm. Neuronal activity was recorded at a 15 Hz frame rate (796 x 512 pixels) for 10 s per trial by detecting fluorescence changes using a photomultiplier tube (Hamamatsu). Imaging was intermitted during the 2s ITI.

#### $Ca^{2+}$ imaging analysis

$Ca^{2+}$ imaging data of each session was processed using the Python-based Suite2p processing pipeline[78]. Recordings were motion-corrected using a non-rigid registration. Cells were detected using automated ROI detection followed by manual curation to ensure no false labeling. Raw fluorescence time courses were extracted as each ROI's (non-weighted) mean pixel value. Additionally, a neuropil trace was computed for each ROI to generate neuropil-corrected calcium traces (F(t), t = time). Change of fluorescence was calculated by subtracting the baseline fluorescence $F_0$ (mean fluorescence value of the first 1.5 s) from the corrected traces F(t) and dividing by $F_0$:

$$\frac{\Delta F}{F} = \frac{\mathrm{F(t)} - \mathrm{F_0}}{\mathrm{F_0}}$$

#### Alignment of cell masks across days

Individual sessions recorded on different days were aligned using the MATLAB-based ROIMatchPub package (available at https://github.com/ransona/ROIMatchPub/tree/master) pipeline. All sessions were registered, and several reference ROIs were manually labeled across sessions. ROIs of all sessions were then automatically aligned and mapped, enabling longitudinal tracking of the same neurons across all imaging sessions. Only neurons that were detected across relevant sessions were used for further analyses, including tensor component analysis (TCA), temporal decoding, selectivity index (SI) and history modulation index (HMI).

#### Criteria for active neurons

For tensor component analysis and subsequent decoding of behavioral strategies, only longitudinally recorded neurons tracked across all sessions were included. Further analyses, such as calculation of SI and HMI (in the respective session), as well as functional classification of neurons (in both respective pre- and post-reversal sessions), were only conducted for neurons that met the following activity ($\Delta F/F$) criteria:

- Peak response (for stimulus or reward-outcome window) is at least a 25% increase.

- Response (during stimulus or reward-outcome window) significantly ($p < 0.05$, t-test) different from average baseline response (1.5 s before the texture was presented).
- Mean response (for stimulus-presentation or reward-window) more than 3 × noise from the baseline.

**Selectivity index**

We calculated the selectivity of individual neurons for a specific trial type over another using a receiver operating characteristic (ROC) analysis as previously described[9]. This was used to quantify the ability of an ideal observer to discriminate between said trial types based on single-trial responses[75,79]. Based upon the possible outcomes of using a strategy in question, we assessed selectivity either for 'hit' against 'miss' ('hit-stay cue') or for 'FA' against 'CR' ('FA-shift cue') trials. ROC analysis was conducted on the 1.5 s segment of the $\Delta F/F$ transients during the outcome window. Based on the dot-product similarity of the transient to the mean transient for the same trial type minus the dot-product similarity to the mean for the other trial type, a 'discrimination variable' score (DV) was determined for each trial. We therefore computed for 'hit' trials,

$$\mathrm{DV_{hit}} = H_i(\bar{H}_{\forall j \neq i} - \bar{M})$$

for 'miss' trials,

$$\mathrm{DV_{miss}} = M_i\left(\bar{M}_{\forall j \neq i} - \bar{H}\right)$$

for 'FA' trials,

$$\mathrm{DV_{FA}} = F_i\left(\bar{F}_{\forall j \neq i} - \bar{C}\right)$$

and for 'CR' trials,

$$\mathrm{DV_{CR}} = C_i(\bar{C}_{\forall j \neq i} - \bar{F})$$

where $H_i$, $M_i$, $F_i$, $C_i$ are the calcium transients of the $i$-th hit, miss, FA, CR trial under consideration respectively, and $\bar{H}$, $\bar{M}$, $\bar{F}$, $\bar{C}$ are the mean transients. The 'selectivity index', SI, was determined by calculating:

$$\mathrm{SI} = 2 \times (\mathrm{AUC} - 0.5)$$

Where AUC is the area under the curve for a binary classifier of the two outcomes in question after being trained on the discrimination variable. Negative SI values indicate a preference for the non-responsive outcomes (preferring ‘miss’ over ‘hit’ or preferring ‘CR’ over ‘FA’ respectively) while positive SI values indicate a preference for the responsive outcomes (preferring ‘hit’ over ‘miss’ or preferring ‘FA’ over ‘CR’ respectively). Statistical significance ($p < 0.05$) was tested by comparing SIs to a distribution created by using a permutation test (500 permutations) with randomly shuffled trial labels.

**Functional classification of neurons**

Neurons that met all activity criteria in the pre-reversal period (LE) and at least one of the post-reversal periods (RN or RE) were grouped into functional classes based on the change of their SI value upon rule-switch as described previously[9]. If the two SI values before and after reversal, respectively, were of the same sign and met the significance criterion, a neuron response was classified as ‘outcome-selective’. This means such a neuron had a consistently higher amplitude for one of the two compared outcomes, before and after reversal, independent of stimulus identity. Neurons classed as ‘outcome-selective’ were used for further analyses, including strategy decoding and history-modulation index. ‘Stimulus-selective’ neurons were classed as such if they responded to ‘hit’ before reversal but ‘FA’ after reversal (or vice versa), indicating they were responding to the stimulus presented (P100 or P1200) rather than the outcome.

**Tensor component analysis**

Calcium traces from imaged S1 neurons for each mouse were organized into one tensor $X$ per mouse, where $X_{nkt}$ denotes the nonnegative activity of neuron $n$ at time $k$ on trial $t$

normalized ([0-1]) for each neuron separately. $X$ has dimensions N x K x T where N is the total number of neurons, K is the total imaged frames in a trial and T is the total number of trials. Tensor component analysis (TCA)[30] was carried out on tensors separately (separate decompositions for each mouse) using the Nonnegative Matrix and Tensor Factorization Toolbox in MATLAB (available at https://github.com/kimjingu/nonnegfac-matlab). The nonnegative CP decomposition of all tensors was carried out using the Hierarchical Alternating Least Squares (HALS) method[30]. To determine the component number (rank) for each decomposition, the reconstruction error and similarity of solutions were tested for components 1-10/11. The target rank that achieved the lowest reconstruction error whilst maintaining a sufficient similarity between solutions (>0.7) was selected as the target rank. However, when similarity achieved very high levels (>0.95) for components with similar reconstruction error (1-2 components away), this component was selected instead. For mice that were lOFC silenced, a single session from the LE, RN and RE phases were taken and concatenated into a tensor $X_{silenced}$ for each silenced mouse separately and decomposed into components. To create an appropriate comparison of WT TCA components to the silenced mice, $X_{WT\ compare}$ was created, which concatenates only a single session from the LE, RN and RE phases before being decomposed. FA-CR 'outcome-selective' neurons were identified using the functional classification of neurons mentioned in the previous section. These 'outcome-selective' neurons responded selectively to FA-CR in the LE and RE task phases. Tensor decomposition was also run on only these select neurons from $X_{silenced}$ and $X_{WT\ compare}$, as well as a population-matched non-outcome selective neuron control group. This pipeline was also completed for 'stimulus-selective neurons'.

**Component classification**

To investigate whether components exhibited selective responses to sandpaper texture (stimulus), licking (action), hit/FA (outcome) or other task features, we grouped components into distinct types based on their time factor. We calculated the time points at which a component has a positive onset of activity (activity beginning to increase, defined at the timepoint of derivative change prior to a significant activity increase calculated in a sliding window averaging over three datapoints on each side) or a negative onset of activity (activity beginning to decrease, defined as above but prior to a significant activity decrease) within a trial. If a component contained a single onset (positive or negative) that occurred in the stimulus, action or outcome windows, they were labelled a ‘stimulus’, ‘action’ or ‘outcome’ component type, respectively. If a component contained an onset in none of the stimulus, action, or outcome windows, it was labelled as an ‘other’ type. If components showed multiplexed activity (multiple onsets in different windows), they were labelled as follows: If a component had one onset inside either the ‘sensory’, ‘action’ or ‘outcome’ window, and another onset outside of these windows, the component was labelled with the onset outside of the task-relevant windows ignored. If a component had two or more onsets inside the ‘sensory’, ‘action’ or ‘outcome’ windows, separately, they were labelled as ‘other’. For example, if a component had onsets in the ‘action’ window and in the ‘outcome’ window, this component would be classified as an ‘other’ type component. After classifying a component by its time factor, we used the associated task variable, based on the component’s type, to group its trial factors. For example, if a component was classed as a ‘stimulus’ type we separately grouped the trial factors values, where P100 (stimulus 1) was presented, together and grouped the trial factors, where P1200 (stimulus 2) was presented, together for that component. The same was done for ‘action’ and ‘outcome’ component types.

**Component demixing**

To extract a measure for the selectivity of a component for a specific task variable, we defined ‘component demixing’. ‘Component demixing’ quantifies the absolute separation between normalized ([0-1]) trial factors belonging to the relevant task variables for that component type. For example, for a ‘stimulus’ component type, component demixing will quantify the average scaled separation between the trial factors of P100 trials and P1200 trials. This was done separately within each of the four task phases respectively (LN, LE, RN, and RE).

**Strategy decoding analysis**

To decode strategy representations within S1, we developed a novel strategy decoding pipeline. For each mouse, trial factors from all TCA components were concatenated into a matrix (components X trial factors). Next, for each chosen strategy we defined the strategy vector, S(*t*), indicating if P(strategy) was increased (strategy used, S(*t*) = 1), remained constant (no evidence, S(*t*) = 0), or decreased (strategy avoided, S(*t*) = -1) for a given trial *t*. Strategy decoding analysis was performed by training ('training trials') and subsequent evaluation ('test trials') of a multinomial logistic regression (MLR) classifier (*mnrfit* function MATLAB) for decoding strategy usage, S(*t*), in a sliding trial window. More technically, the average classification accuracy across 200 iterations (Monte-Carlo cross-validation) for a sliding trial window with discrete steps was extracted.

Strategy decoding was performed at either 'fine' or 'coarse' trial resolutions. For 'fine' trial resolution temporal decoding analysis, trials were binned into a sliding window of length 210 trials and a separation of 100 trials between windows. Window lengths were chosen to ensure a sufficient number of strategy use or avoidance trials were present. The sliding window was adjusted to never include trials from both the learning and reversal phases. For a given window, 90% of trials were used for training, and 10% of held-out trials were used to evaluate the MLR classifier. For the 'training trials', trial factor values from all TCA components with their corresponding strategy use vector S('training trials') were used to train the classifier. The training dataset was subsampled so that the number of the most frequent and second most frequent categories (employ, no evidence, avoided) were equal in order to avoid biasing the classifier towards the most populous category. The trained classifier then estimated probabilities for each possible value of S(*t*) (employed, no evidence, avoided) based on the 'test trials' trial factor values from all components. We used a 'winner takes all' method such that the event with the highest probability allocated to it by the classifier was taken as the classifier's prediction of strategy response: S('test trials')$_{\text{predicted}}$. The predicted strategy events were then compared to the true strategy updates to compute the classification accuracy. The following equations formalize these steps

$$A_{I,W} = \frac{\sum_{\mathrm{k=test\ trials}_{I,W}} \delta_{\mathrm{S(k)_{predicted}},\mathrm{S(k)}}}{\mathrm{N}_{\mathrm{test\ trials}_{I,W}}} \qquad \mathrm{Accuracy}(W) = \frac{\sum_{I=1}^{200} \mathrm{A}_{I,W}}{200} \times 100$$

Where 'test trials$_{I,W}$' are the tested trials for iteration $I$ and trial window $W$. $\mathrm{N}_{\mathrm{test\ trials}\ I,W}$ is the number of test trials (equal to 21 for 'fine' resolution). $\mathrm{A}_{I,W}$ is the proportion of correct predictions of the classifier in iteration $I$ and trial window $W$. Accuracy($W$) is the decoding accuracy (%) of the classifier for a given strategy in window $W$. $\delta_{xy}$ is the Kronecker delta function. To compare the decoding accuracy across the four task phases (LN, LE, RN, RE), we used a 'coarse' trial resolution strategy decoding analysis where 250 trials were chosen from each task phase to be the trial window. Subsampling of strategy use, avoidance and no-evidence trials were used to match trial numbers across categories to avoid biasing the model where possible. If insufficient trials were found at any iteration for the software, additional trials were added to the subsampled training data.

To evaluate the impact of outcome component types on strategy decoding, we held out trial factors from outcome components and reran strategy decoding. Strategy decoding for lOFC-silenced mice trial factors was implemented at 'coarse' resolution and compared to strategy decoding of WT mice. For this analysis trial factors of WT mice were taken from the Tensor decomposition of the $X_{WT\ compare}$ tensor, while trial factors of lOFC silenced mice were taken from $X_{silenced}$. Strategy decoding from FA-CR outcome selective cells and population matched controls were found by applying decoding on the separate tensor decompositions for these two neuron groups.

To confirm that strategy decoding could not be explained by only S1 population codes of outcome, we trained a separate MLR outcome classifier to identify outcomes that could inform a 'guess' of the strategy used: 1) 'hit', 'miss' and 'other' outcome categories were used for the outcome classifier to compare to strategy classification of 'hit-stay cue', as these are the possible outcomes of 'use', 'avoidance' or 'no evidence'. 2) 'hit', 'miss', 'CR', 'FA' for 'hit-stay choice' 3) 'FA', 'CR' and 'other' for 'FA-shift cue' 4) 'FA', 'CR', 'hit', 'miss' for 'FA-shift choice'. For 'hit-stay cue', at each iteration we extracted the outcome classifier's prediction (winner-takes all) of 'hit', 'miss' and 'other' within the test trials. We then replaced each trial of the outcome classifier's predicted 'hit' trials with 'strategy-guesses' of 'use', 'avoid', and 'no evidence' events. Each 'strategy-guess' for each 'hit' trial was done with a 'weighted' probability, where the probability of a given predicted 'hit' trial being replaced by 'use' was the same as the proportion of predicted 'hit' trials that were also predicted as 'use' by the MLR strategy-classifier, and so on for 'avoid' and 'no-evidence'. The same process defined the strategy-guess for the outcome classifier's predicted 'miss' and predicted 'other' trials, thus building strategy guesses at each iteration based on outcome decoding. The same protocol was also carried out for 'hit-stay choice', 'FA-shift cue' and 'FA-shift choice' for their relevant outcomes.

**History modulation index**

To assess the influence of previous outcome experience on neural activity during the 1.5 s outcome window on subsequent trials, we calculated strategy-specific history modulation indices (HMI). For 'hit-stay cue' we calculated how the response magnitude during a 'hit' trial was altered, depending on if the previous trial was rewarded ($R_{hit-hit}$) or non-rewarded ($R_{nohit-hit}$) (Reward-HMI). To perform this calculation, we normalized the difference between 'hit' trial responses preceded by a reward ('hit') compared to any other outcome ('nohit'):

$$\mathrm{RHMI} = \frac{\mathrm{R_{nohit-hit}} - \mathrm{R_{hit-hit}}}{\mathrm{R_{hit}}}$$

For 'FA-shift cue' we calculated how the response magnitude during a 'CR' trial was altered, depending on whether the previous trial was rewarded ($R_{FA-FA}$) or not punished ($R_{noFA-FA}$) (Error-HMI). To perform this calculation, we normalized the difference between 'CR' trial responses preceded by a punishment ('FA') compared to any other outcome ('noFA'):

$$\mathrm{EHMI} = \frac{\mathrm{R_{noFA-CR}} - \mathrm{R_{FA-CR}}}{\mathrm{R_{CR}}}$$

The respective task phases for which HMIs were calculated are described in the figure legends and main text.

**Statistical analysis**

The statistical analyses used in this study are described in the respective figure legends and the main text. Non-parametric tests such as two-sided Wilcoxon rank-sum tests and permutation tests were used unless otherwise stated, allowing us to avoid making excessive assumptions about data distributions. Student’s t-test was used 1) when sample sizes were sufficient ($n>30$) and data distributions passed the Kolmogorov-Smirnov test, or 2) when previous literature or small datasets supported doing so. Predetermined sample sizes were not found using statistical methods. Assessments of outcome and allocation during experiments were not made blind to investigators.

### Supplementary Figure 1

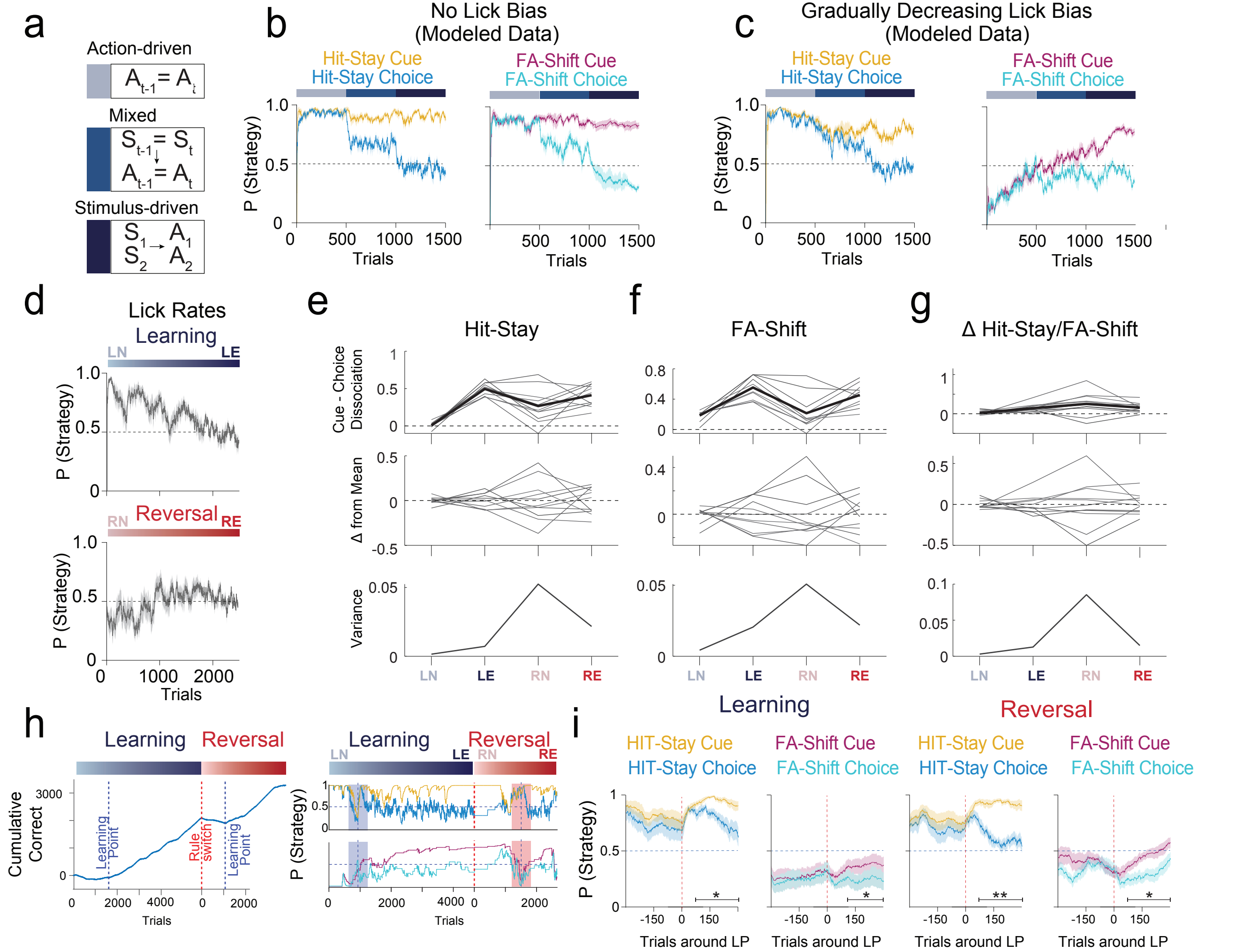


**Supplementary Figure 1. The highest individual variability of strategies occurs immediately after reversal.** **a**, Schematic of the three different states of modeled agents, following a purely action-driven, stimulus-driven or mixed strategy. **b**, Comparison of cue- and choice-based 'hit-stay' and 'FA-shift' strategies displayed by modeled agents during the three respective task states. **c**, same as **b**, but with a gradually decreasing lick bias incorporated. **d**, Lick rates of animals displaying a high lick bias during the naïve learning phase, which gradually decrease with training. **e**, Different measurements of individual variability in reward-driven strategy use ('hit-stay') across salient task phases (LN, LE, RN, RE). Top, averaged (black) and individual (grey) cue-choice dissociation. Middle, the difference of each individual animal from the average cue-choice dissociation. Variance across all animals. **f**, Same as **e** but for error-driven learning ('FA-shift'). **g**, Same as **e** but for the difference of cue-choice dissociation between strategies ('hit-stay' cue-choice dissociation - 'FA-shift' cue-choice dissociation). **h**, Schematic of learning points for one example mouse. Learning points are defined when the derivative of the curve displaying cumulative correct decisions becomes positive. **i**, Comparison of cue and choice-based strategy dynamics for 'hit-stay' and 'FA-shift' strategies around the learning points during learning and reversal phases. *n* = 11 WT mice for all plots. Data presented as mean ± S.E.M., horizontal bars indicate significant differences between curves at the respective trials, cluster-based permutation test.

### Supplementary Figure 2

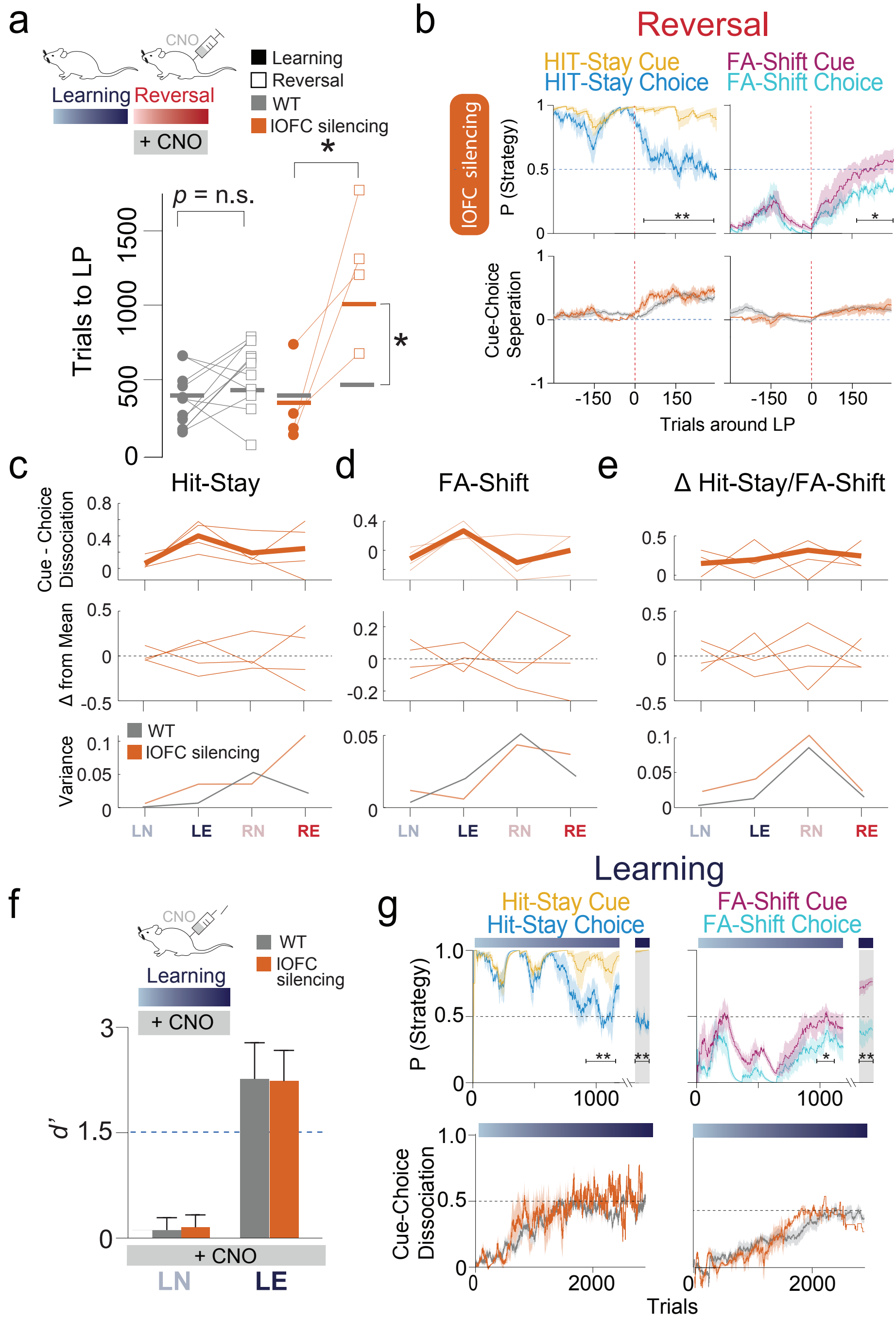


**Supplementary Figure 2. lOFC-silencing affects individual variability only for reward-driven adaptation after reversal. a**, Comparison of trial numbers between animals with lOFC-silencing during the reversal phase and WT controls needed to reach the learning point during reversal learning. **b**, Top, comparison of cue and choice-based strategy dynamics for 'hit-stay' and 'FA-shift' strategies in lOFC-silenced animals (silenced during reversal) around the learning points during learning and reversal phases. Bottom, comparison of the cue-choice dissociation for the respective strategies for both groups during learning and reversal phases. **c**, Different measurements of individual variability in reward-driven strategy use ('hit-stay') across salient task phases in lOFC-silenced mice (LN, LE, RN, RE). Top, averaged (bold orange) and individual (orange) cue-choice dissociation. Middle, the difference of each individual animal from the average cue-choice dissociation. Variance across all animals. **d**, Same as **c** but for error-driven learning ('FA-shift'). **e**, Same as **c**, but for the difference of cue-choice dissociation between strategies ('hit-stay' cue-choice dissociation - 'FA-shift' cue-choice dissociation). **f**, Comparison of behavioral performance (*d*') of lOFC-silenced mice during initial learning and WT controls aligned to the session number. WT average reached the respective task phase. Neuronal silencing was achieved by daily systemic injection of CNO during initial learning (LN→LE). **g**, Comparison of cue and choice-based strategy dynamics from lOFC-silenced mice during learning for 'hit-stay' and 'FA-shift' strategies. *n* = 4 lOFC-silenced mice and *n* = 11 WT mice for all plots. Data presented as mean ± S.E.M., *p<0.05, two-tailed t-test. Kolmogorov-Smirnov test. Horizontal bars indicate significant differences between curves at the respective trials, cluster-based permutation test.

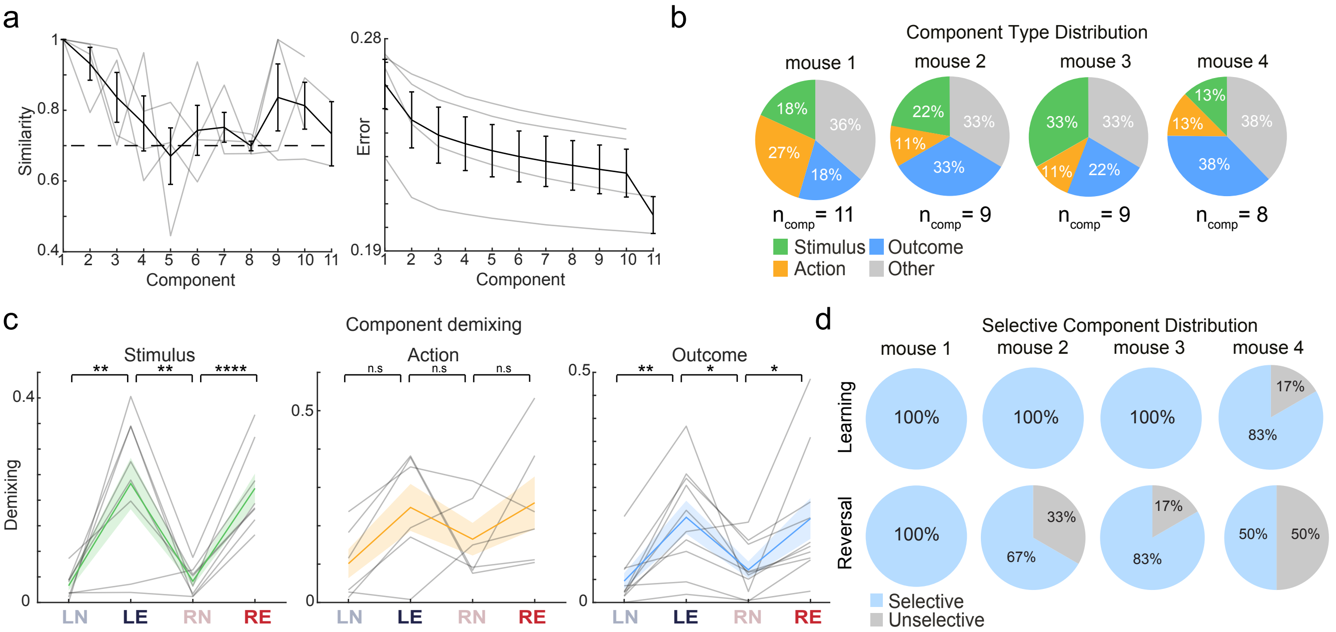


**Supplementary Figure 3. Stimulus, action and outcome representations in the S1 neuron population. a**, Left, individual (grey) and mean (black) similarity scores for different TCA component numbers. The dotted line indicates the minimum threshold, 0.7, for the chosen number of components (*n* = 4 mice, 836 active neurons). Right, same as left, but for reconstruction error. **b**, Component type distributions for individual mice. **c**, Component demixing – absolute difference in trial factor activation for the relevant task variable – for all stimulus, action and outcome type components for WT mice in key task phases (LN, LE, RN, and RE). Grey lines show individual components, colored lines show mean component demixing. **d**, Top, selective component distribution for individual WT mice in the learning task phase (LN→LE) and, bottom, in the reversal task phase (RN→RE). Selective components have significantly separated trial factor activation for their corresponding task variable in at least one session. Data displayed as mean ± S.E.M., *p<0.05, **p<0.01, ****p<0.0001, Kolmogorov-Smirnov test, two-sided t-test, two-sided Wilcoxon rank-sum test, permutation test, Bonferroni-Holm correction for multiple comparisons.

### Supplementary Figure 4

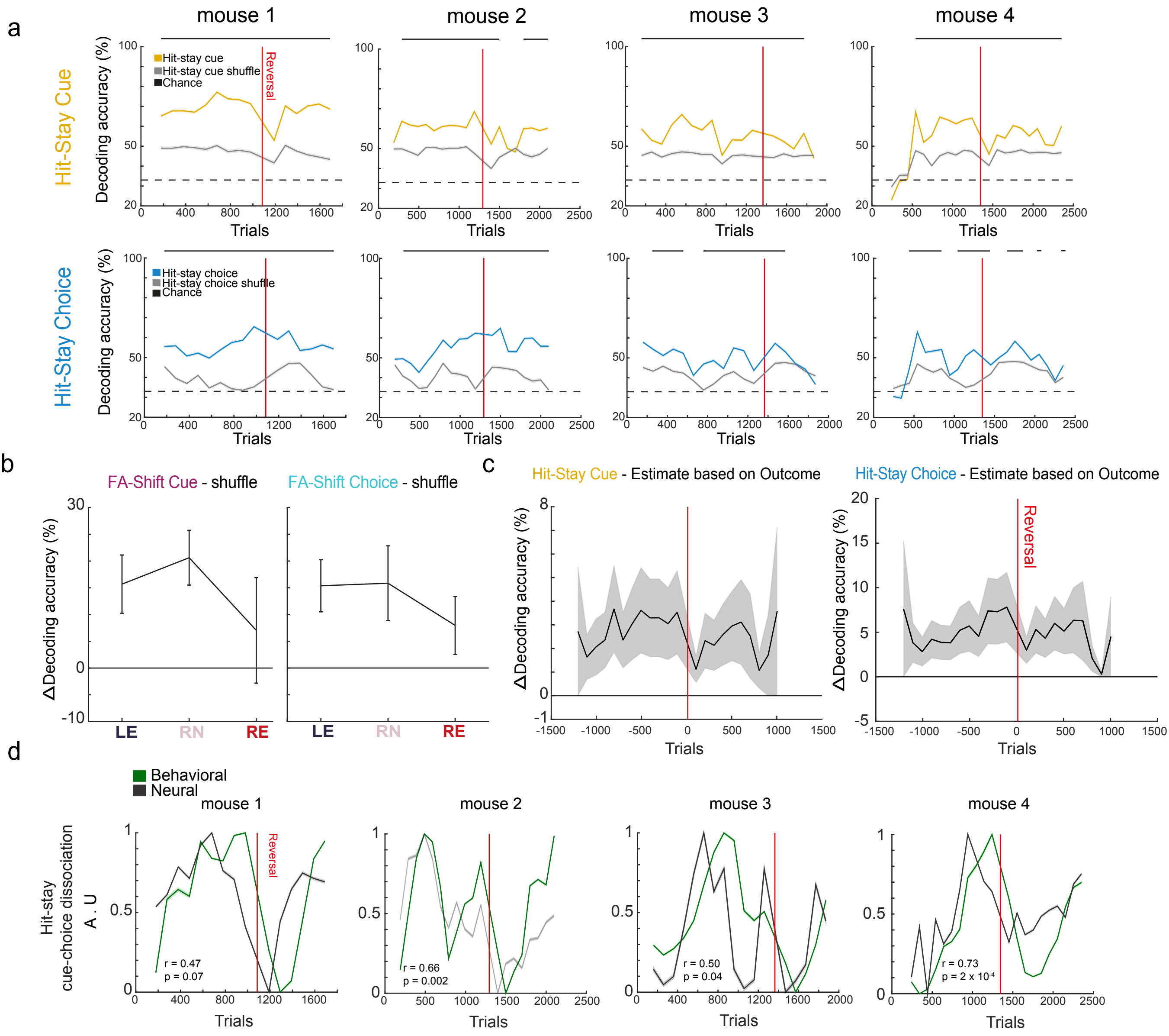


**Supplementary Figure 4. S1 neural population representations of strategy are stable across mice. a**, Strategy-decoding plots for 'hit-stay' cue and choice for unshuffled and shuffled S1 trial factor data for individual WT mice (*n* = 4 mice, 836 neurons). 'Hit-stay' classification accuracy (Monte Carlo cross-validation, 200 iterations, see Methods for detailed description) for all cue and choice strategies significantly exceeded classification accuracy from shuffled trial factors for almost all trial windows. Chance-level decoding (33%) is shown by dotted lines. **b**, Left, difference between decoding accuracy of 'FA-shift cue' for unshuffled and shuffled trial factors in key task phases (LE, RN, and RE). Right, same for choice strategy. 'FA-shift' decoding accuracy for cue and choice strategies significantly exceeded decoding accuracy of shuffled data over the task ($p<0.01$, permutation test). **c**, Left, difference between decoding accuracy of 'hit-stay cue' and the best estimate of 'hit-stay cue' assuming only outcome encoding in S1 is used for strategy classification (Accuracy('hit-stay cue') - Accuracy('hit-stay cue' estimated)) (see Methods for detailed description). Right, same as left, but for 'hit-stay choice'. 'Hit-stay' decoding accuracy for all cue and choice strategies significantly exceeded decoding accuracy from estimated strategies based on outcome over the task ($p<0.0001$, permutation test). **d**, Normalized behavioral cue-choice dissociation (P(cue) - P(choice)) and the respective normalized dissociation of decoding accuracy (Accuracy(cue) - Accuracy (choice)) for 'hit-stay' for individual mice. r, correlation coefficient; A.U, arbitrary units. Data displayed as mean ± S.E.M., horizontal bars above plots indicate significant ($p<0.05$), permutation test, Bonferroni-Holm correction for multiple comparisons, Pearson correlation coefficient and associated t-statistic.

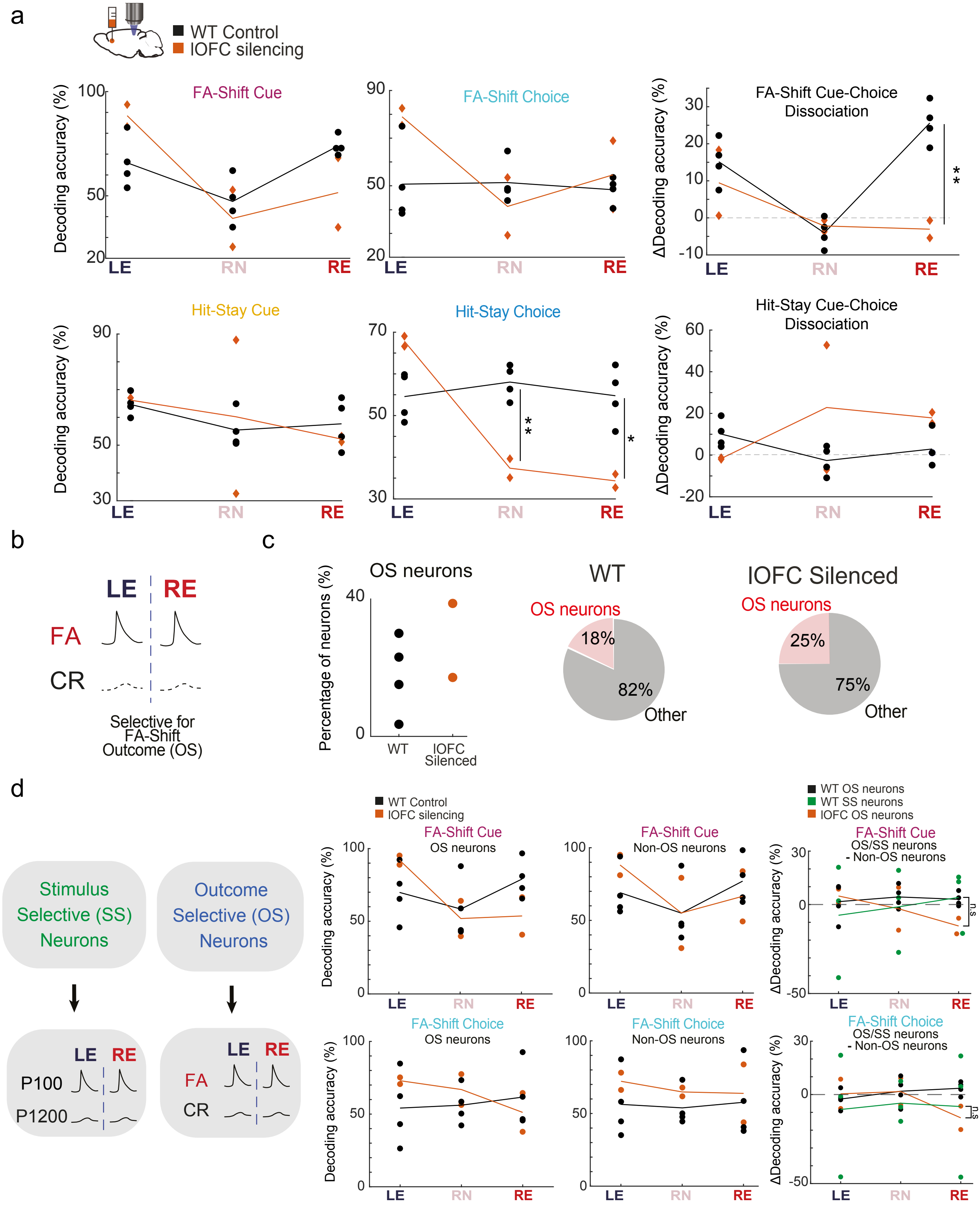


**Supplementary Figure 5. lOFC-silencing disrupts post-reversal strategy representations in the S1 neuron population.** **a**, Top row, decoding accuracy from S1 neuron population trial factors of 'FA-shift cue', 'FA-shift choice' and their dissociation (Accuracy(cue) - Accuracy(choice)) in key task phases (LE, RN, and RE) for WT ($n$ = 4 mice) and lOFC-silenced mice that were silenced during the reversal phase ($n$ = 2 mice). Bottom row, same as top, but for 'hit-stay'. **b**, Schematic of outcome-selective (OS) neurons that respond to 'FA-shift cue' relevant outcomes (FA, CR). **c**, Proportion of FA-CR OS neurons as a fraction of all active neurons from WT mice and lOFC-silenced mice is shown, for individual mice (scatter plot) and all mice (pie charts). **d**, Left, schematic of outcome- and stimulus-selective (SS) neuron definition. Right, top row, decoding accuracy of 'FA-shift cue' from OS neuron trial factors, non-OS neuron trial factors (neuron numbers matched), and their difference (OS – Non-OS), as well as the difference for SS neurons (SS – Non-OS), is shown before (LE) and after reversal (RN, RE). Right, bottom row, same as top row, but for 'FA-shift choice'. Scattered points indicate individual mice, lines indicate the mean. *p<0.05, **p<0.01, Kolmogorov-Smirnov test, two-sided t-test.

### Supplementary Figure 6

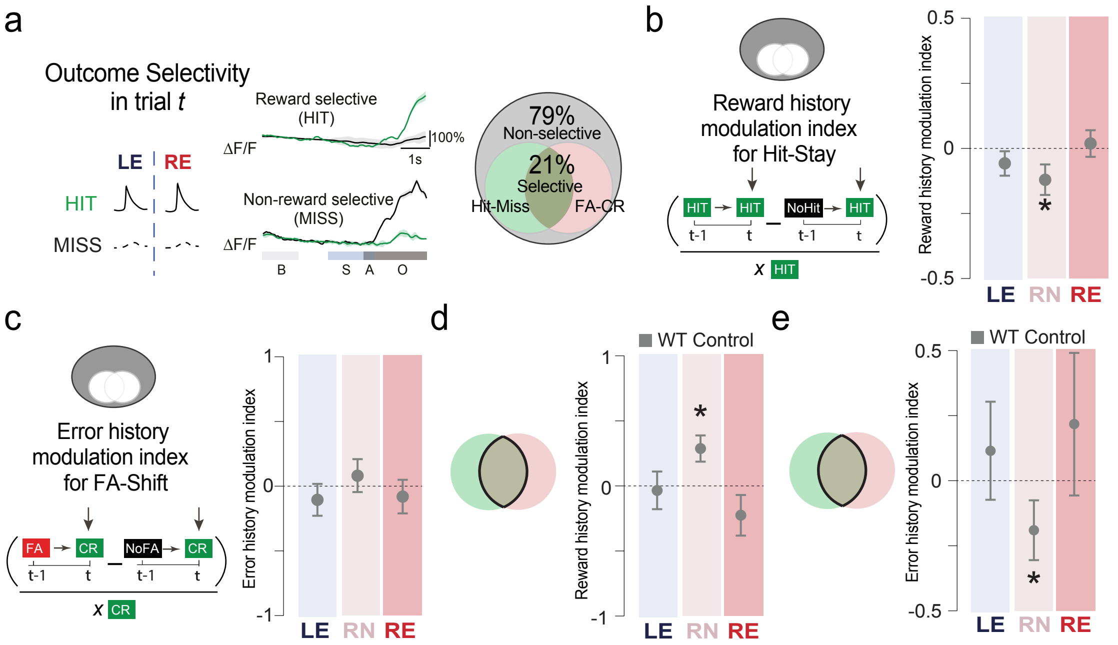


**Supplementary Figure 6. Non-selective and mixed selective cells show minimal history modulation after reversal. a**, Left, schematic and neural responses of 'hit-selective' neurons. Right, proportion of strategy-specific 'outcome-selective' neurons in the total cell population. 'Non-outcome-selective' neurons are displayed in grey. **b**, Left, calculation of the 'hit-stay cue' associated reward history modulation index (RHMI). Neural responses of 'hit' trials (defining strategy use in trial *t*), preceded by a 'hit' trial, and those preceded by a 'non-hit' trial are subtracted and divided by the average 'hit' trial response to indicate reward-driven history modulation (positive values) for 'hit-stay' strategy-specific variables. Right, RHMI for non-outcome-selective neurons in WT animals across key task phases (LE, RN, and RE). **c**, Left, calculation of the 'FA-shift cue' associated error modulation index (EHMI). Neural responses of 'CR' trials (defining strategy use in trial *t*) preceded by a 'FA' trial and those preceded by a 'non-FA' trial are subtracted and divided by the average 'CR' trial response to indicate error-driven history modulation (positive values) for 'FA-shift cue' strategy-specific variables. Right, EHMI for non-outcome-selective neurons in WT animals across task phases (LE, RN, and RE). **d**, RHMI for 'mixed-selective' neurons. **e**, EHMI for 'mixed-selective' neurons. *n* = 3 WT mice. Data displayed as mean ± S.E.M., *p<0.05, two-tailed t-test.
